# Nonspecific membrane bilayer perturbations by ivermectin underlie SARS-CoV-2 *in vitro* activity

**DOI:** 10.1101/2023.10.23.563088

**Authors:** Richard T. Eastman, Radda Rusinova, Karl F. Herold, Xi-Ping Huang, Patricia Dranchak, Ty C. Voss, Sandeep Rana, Jonathan H. Shrimp, Alex D. White, Hugh C. Hemmings, Bryan L. Roth, James Inglese, Olaf S. Andersen, Jayme L. Dahlin

## Abstract

Since it was proposed as a potential host-directed antiviral agent for SARS-CoV-2, the antiparasitic drug ivermectin has been investigated thoroughly in clinical trials, which have provided insufficient support for its clinical efficacy. To examine the potential for ivermectin to be repurposed as an antiviral agent, we therefore undertook a series of preclinical studies. Consistent with early reports, ivermectin decreased SARS-CoV-2 viral burden in *in vitro* models at low micromolar concentrations, five-to ten-fold higher than the reported toxic clinical concentration. At similar concentrations, ivermectin also decreased cell viability and increased biomarkers of cytotoxicity and apoptosis. Further mechanistic and profiling studies revealed that ivermectin nonspecifically perturbs membrane bilayers at the same concentrations where it decreases the SARS-CoV-2 viral burden, resulting in nonspecific modulation of membrane-based targets such as G-protein coupled receptors and ion channels. These results suggest that a primary molecular mechanism for the *in vitro* antiviral activity of ivermectin may be nonspecific membrane perturbation, indicating that ivermectin is unlikely to be translatable into a safe and effective antiviral agent. These results and experimental workflow provide a useful paradigm for performing preclinical studies on (pandemic-related) drug repurposing candidates.

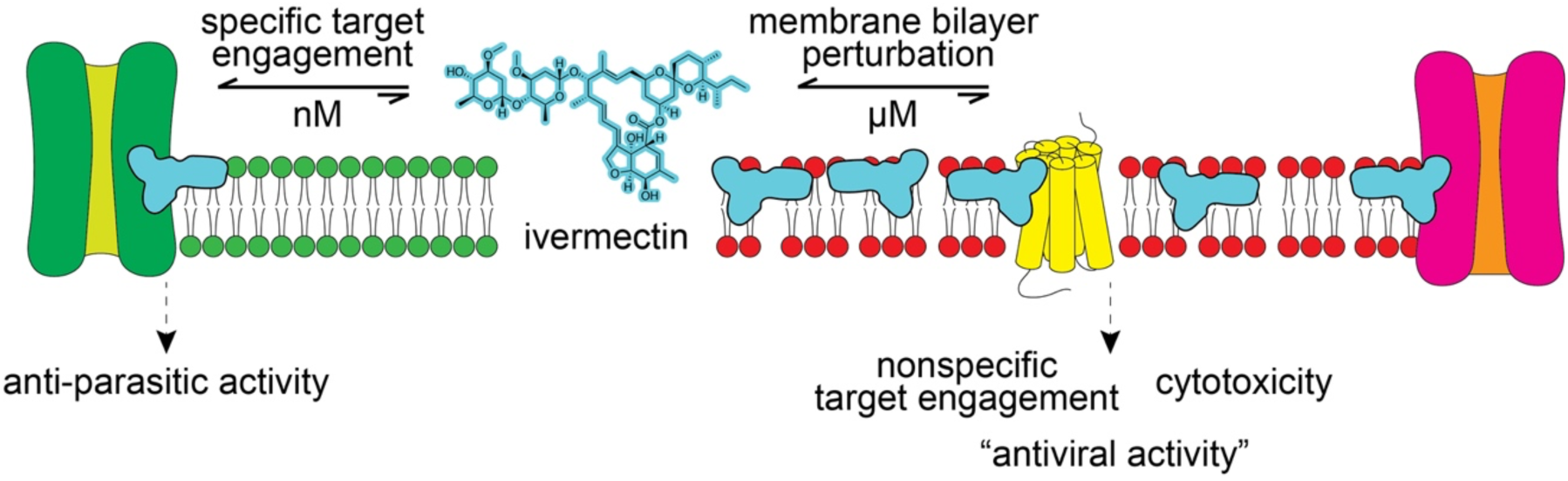
Graphical abstract.

## Introduction

Ivermectin (IVM) is an important antiparasitic drug for diseases such as onchocerciasis, strongyloidiasis, ascariasis, and lymphatic filariasis.^1^ IVM is a hydrophobic molecule that inserts into the outer leaflet of plasma membranes, where it binds at the subunit interfaces of transmembrane protein domains such as Cys-loop receptors.^2^ IVM kills parasites by activating a glutamate-gated Cl-channel which causes parasite muscle and nerve hyperpolarization with subsequent paralysis and death. Importantly, IVM does not appear to cross the plasma membrane and binds to these classical targets at *low nanomolar* concentrations.^3, 4, 5, 6^ IVM also alters the function of other transmembrane targets including the farnesoid X receptor, the human *ether-‘a-go-go*-related gene (“hERG”) potassium channel, and G-protein-gated inwardly rectifying potassium channel (**Supplementary Table 1**).^2^

It is important to consider the natural history of IVM as a reported antiviral agent. IVM was originally proposed as an antiviral agent based on a biochemical (cell-free) AlphaScreen to identify small-molecule nuclear import inhibitors.^7^ In that report, the target was the HIV integrase (a nuclear localization signal, NLS) and mouse importin α-β1 protein-protein interaction. Years later, in the early phases of the Covid-19 pandemic, IVM was reported to have antiviral activity at low micromolar concentrations by decreasing SARS-CoV-2 RNA in Vero/hSLAM cells without any reported cytotoxicity.^8^ Follow-up biophysical studies at high micromolar concentrations (∼25 to 100 μM) demonstrated that IVM could disrupt its purported antiviral target, the host importin α/β1 heterodimer.^9^ These results, however, should be interpreted in the context that the estimated comatose/fatal plasma concentration of IVM (total drug) is about 100 nM and that the solubility limit for IVM is 1 to 2 µM.^10, 11, 12^ Moreover, in *Ascaris suum* muscle cells, IVM is primarily localized to the *outer* leaflet of the plasma membrane and is not found in organellar membranes, suggesting it has relatively low intracellular concentrations.^6, 13^ Along with its effect on SARS-CoV-2, IVM was reported to have antiviral activity against numerous related and unrelated viruses generally at low micromolar concentrations.^9, 14^

We evaluated the reported SARS-CoV-2 antiviral activity of IVM in preclinical studies including live-virus assays, cellular health profiling, counter-screens for assay interferences, and target selectivity. Whereas IVM showed antiviral activity at low micromolar concentrations, in agreement with original reports, this activity closely paralleled multiple biomarkers of cellular injury. IVM was likewise found to perturb membrane bilayers at low micromolar concentrations, where it can nonspecifically modulate membrane-based targets (**Supplementary Table 1**). These data provide a molecular mechanism to explain the observed lack of translatability of IVM as a SARS-CoV-2 therapeutic and provide important caveats regarding drug repurposing.

## Results

### SARS-CoV-2 antiviral activity *in vitro*

IVM was tested for antiviral activity in a live-virus assay in A549-ACE2 cells using SARS-CoV-2 labelled with an FLuc reporter.^15^ Three independent samples of IVM (**1a**-**c**) showed similar antiviral activity after 48 h compound treatment in this FLuc-based viral assay (mean half-maximal effective concentration, or IC_50_ = 6.8 μM; **Figure 1A**). Given the reported low micromolar aqueous solubility of IVM^10, 11^, all IVM concentrations reported in this study are nominal concentrations (total added amount per volume). This activity occurred at similar IVM concentrations in a toxicity counter-screen measuring host cellular ATP content (CellTiter-Glo; mean half-maximal cytotoxic concentration, or CC_50_ = 10.8 μM; **Figure 1A**). Remdesivir (**16**), an FDA-approved SARS-CoV-2 RNA-dependent RNA polymerase (RdRp) inhibitor used as a positive control, decreased viral burden without gross decreases in cellular viability (IC_50_ = 1.3 μM, CC_50_ > 50 μM; **Figure 1A**). The low micromolar antiviral potency and the lack of accompanying cytotoxicity of remdesivir is generally consistent with previous reports in Vero E6 cells and A549-ACE2 cells^16, 17^, though other reports have also shown higher nanomolar potencies, which may be attributable to differences in cell lines, SARS-CoV-2 strains, and/or assay formats.^18^

**Figure 1.**
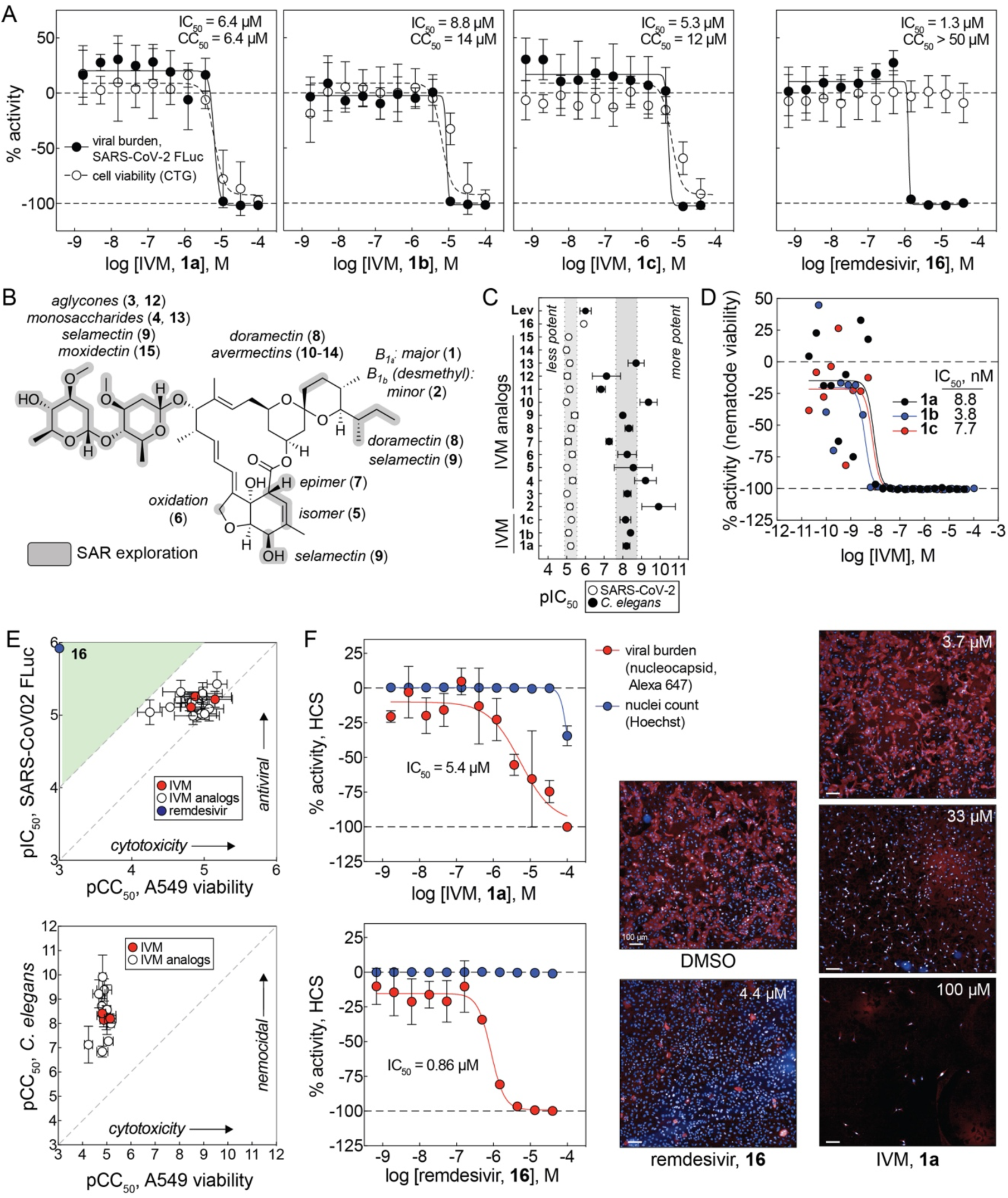
Ivermectin and analogs inhibit SARS-CoV-2 *in vitro* but correlates with host cell toxicity. (A) IVM decreases FLuc-reporter SARS-CoV-2 virus in A549-ACE2 cells at compound concentrations that parallel decreases in host cell viability. IVM was tested with three samples each from different vendors (**1a**-**c**). Remdesivir (**16**), a positive non-toxic antiviral control compound, decreases FLuc-labelled SARS-CoV-2 virus without affecting cell viability. Data are mean ± SD of five (SARS-CoV-2 FLuc) experimental replicates or (CTG) four experimental replicates performed in with duplicate technical replicates; stippled lines at 0% denote no effect relative to control; stippled lines at –100% denotes complete cell killing or complete virus reduction. (B) Chemical structure of IVM and analogs tested in the SARS-CoV-2 qHTS assay and cellular viability counter-screen. (C) IVM and analogs **2**-**15** show flat SAR in the SARS-CoV-2 FLuc assay but SAR in a high-throughput *C. elegans* nematode viability assay. Data are mean ± SD of five intra-plate technical replicates (FLuc) or five inter-plate biological replicates (*C. elegans*) with exceptions noted in Source Data. (D) Representative 22-point concentration-response curves of three IVM samples **1a**-**1c**. (E) Top: IVM and analog antiviral activity occurs at concentrations that decrease cell viability. Green, desirable cytotoxicity:activity ratio (CC_50_:IC_50_ > 10). Bottom: IVM and analogs show antinematodal activity across several orders of magnitude in contrast to A549-ACE2 host cell cytotoxicity. (F) IVM (**1a**) decreases SARS-CoV-2 virus in A549-ACE2-TMPRSS2 cells in a high-content orthogonal antiviral assay that quantifies SARS-CoV-2 nucleocapsid protein at similar concentrations as the primary SARS-CoV-2 FLuc assay. Cell health was measured simultaneously using Hoechst staining for nuclei. Remdesivir, positive non-toxic antiviral control compound. Data are mean ± SD of three experimental replicates. Note nuclear count does not overlap viral burden, though nuclear count may not be indicative of cellular health. Shown are representative images. Image scales: 100 µm. Source data are provided as a Source Data file.

A collection of commercially available IVM analogs was also assessed for antiviral activity and cytotoxicity using the aforementioned SARS-CoV-2 FLuc assay and CTG counter-screen, to search for an interpretable structure-activity relationship (SAR). The presence of strong SAR is generally supportive of specific compound-target interactions, and of a compound being readily optimizable by medicinal chemistry.^19^ The IVM analogs **2**-**15** compromised aglycone and monosaccharide forms, isomers, epimers, and analogs with minor structural modifications (**Figure 1B**). Analogs **2**-**15** decreased viral content at a similar low micromolar concentration range as the parent compound IVM, demonstrating a flat SAR (**Figure 1C**). In contrast to the antiviral assay, IVM and analogs displayed SAR spanning approximately three orders of magnitude in a quantitative HTS *C. elegans* assay for nematode viability (**Figures 1C, 1D**).^20^ Like IVM, the antiviral activity of analogs **2**-**15** occurred at the same concentrations where A549-ACE2 cellular viability was reduced in the CTG counter-screen, again with a relatively flat SAR (**Figure 1E**). Of note, there is extensive SAR spanning multiple orders of magnitude for the antiparasitic activity of IVM (**Supplementary Note 1**). The flat SAR pattern in **Figure 1C** would argue against the antiviral and related host cell cytotoxic activities being readily optimizable by medicinal chemistry.

IVM was also tested for antiviral activity in an orthogonal high-content assay quantifying SARS-CoV-2 nucleocapsid protein in A549-ACE2-TMPRSS2 cells. Similar to the SARS-CoV-2 FLuc assay, IVM (**1a**) showed antiviral activity at low micromolar concentrations after 48 h compound treatment, though IVM did not decrease in the number of detected nuclei (IC_50_ = 5.4 μM, CC_50_ > 50 μM; **Figure 1F**). Nuclear counts, however, may not reflect cellular viability, as Hoechst 33342 and other nuclear staining reagents are capable of binding DNA in both viable and dying/dead cells when cells are fixed and permeabilized. The positive control remdesivir decreased viral burden with little effect on cellular viability (IC_50_ = 0.8 μM, CC_50_ > 50 μM; **Figure 1F**). These results indicate that IVM can reliably decrease SARS-CoV-2 content at low micromolar concentrations, though the concentration-response curve for its antiviral activity overlaps with that for cellular injury, meaning that the decrease in SARS-CoV-2 content may be a consequence of events induced from cell trauma.

### Cellular health profiling

IVM was profiled in multiple assays at multiple time points across a broad range of concentrations for its effects on cellular viability under conditions mimicking the primary SARS-CoV-2 FLuc viral assay in A549-ACE2 cells. IVM decreased cellular viability at low micromolar concentrations, as measured by ATP content (CC_50_ = 7.7 μM), cellular reducing capacity with a resazurin substrate (CC_50_ = 8.4 μM), and cellular reducing capacity with a luciferase pro-substrate (CC_50_ = 5.1 μM) after 24 h compound treatment. These decreases in cellular viability by IVM were time– and concentration-dependent, with significant cellular injury occurring between 24 and 72 h, as observed with all three methods (**Figure 2A**).

**Figure 2.**
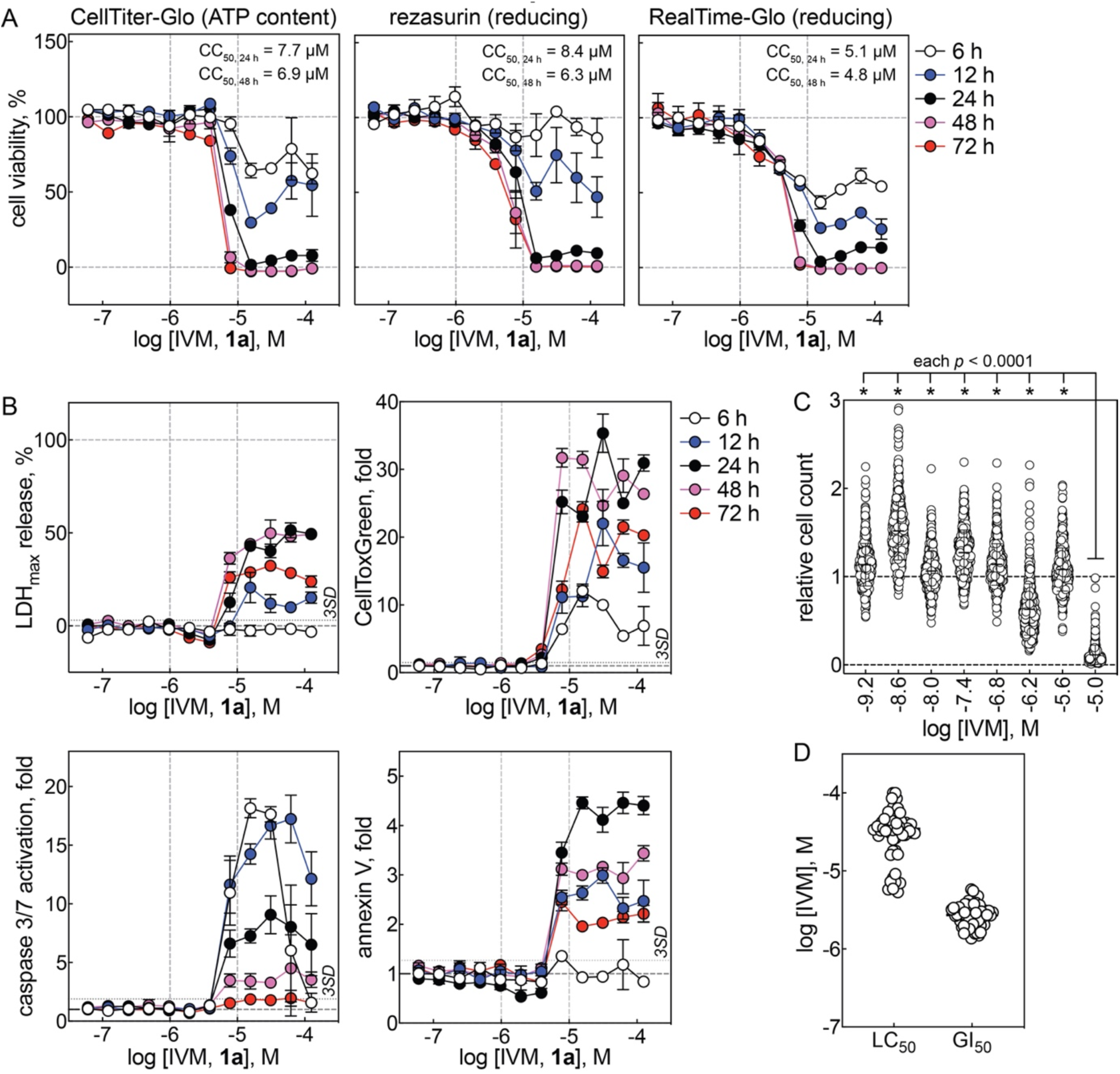
Ivermectin perturbs cellular health at low micromolar concentrations. (A) IVM reduces A549-ACE2 cellular viability at low micromolar concentrations. Cell viability was assayed in three orthogonal formats: CellTiter-Glo (‘CTG’), resazurin, and RealTime-Glo (‘RTG’). Data are mean ± SD of four intra-plate technical replicates; stippled horizontal lines at 100% denote no effect relative to DMSO control; stippled horizontal lines at 0% denotes lack of cell viability relative to digitonin control. (B) Top: IVM causes A549-ACE2 cytotoxicity at low micromolar concentrations. Cytotoxicity was assayed by two orthogonal formats: LDH release and CellTox Green. Bottom: IVM increases biochemical markers of apoptosis in A549-ACE2 cells at low micromolar concentrations. Apoptosis was assayed by caspase 3/7 activation and annexin V binding. Data are mean ± SD of four intra-plate technical replicates; LDH % release calculated relative to digitonin control at time of assay measurment; fold changes calculated realtive to DMSO control for CellToxGreen, caspase 3/7, and annexin V assays. (C) IVM broadly decreases observed cell counts at low micromolar concentrations in the Broad PRISM assay (Broad DepMap, https://depmap.org/portal/). Each data point represents a unique cancer cell line. Statistical differences determined by ordinary one-way ANOVA adjusted for multiple comparisons. (D) IVM broadly decreases observed cell counts at low micromolar concentrations in the NCI60 assay (https://dtp.cancer.gov). Each data point represents a unique cancer cell line. Stippled vertical lines in panels A-C are provided for visual reference. Source data are provided as a Source Data file.

Cellular viability readouts can decrease due to decreases in cellular health and/or decreases in cellular proliferation rates, the cytotoxicity of IVM therefore was also profiled in A549-ACE2 cells, where IVM induced significant (i.e., > 3 SD above baseline) cytotoxicity at low micromolar concentrations (7.8 μM), as measured by lactate dehydrogenase (LDH) release and an orthogonal membrane-impermeable nuclear-binding dye method (CellTox Green). The observed cytotoxicity of IVM was again time– and concentration-dependent, with significant cytotoxicity being observed as early as 6 h, and was similar with both methods (**Figure 2B**). The cytotoxic (non-apoptotic) readout generally peaked before 48 h treatment, with decreased readouts at 72 h, likely due to decreased cellular content.

IVM was similarly profiled for pro-apoptotic effects in A549-ACE2 cells. Treatment with IVM produces massive (> 3 SD above baseline) activation of caspase 3/7 beginning at low micromolar concentrations (7.8 μM) within 6 h. Treatment with IVM also produced significant (> 3 SD above baseline) increases in annexin V binding, an orthogonal pro-apoptotic biomarker, beginning at low micromolar concentrations (7.8 μM) within 12 h. These pro-apoptotic effects were also time– and concentration-dependent and similar with both methods (**Figure 2B**). Compared to cytotoxicity, the apoptotic biomarkers generally peaked within 24 h treatment; the decreased readouts at later time points likely are due to decreased cellular content.

The IVM concentrations associated with decreased cellular viability, induction of cytotoxicity, and induction of apoptosis biomarkers closely approximate antiviral efficacy in both the SARS-CoV-2 FLuc assay and the high-content SARS-CoV-2 assay. Staurosporine, a nonspecific kinase inhibitor used as a positive cytotoxic control, was grossly toxic at low nanomolar concentrations (**Supplementary** Figure 1A). By contrast, remdesivir was grossly toxic only at high micromolar concentrations (**Supplementary** Figure 1B). Notably, select membrane bilayer-modifying control compounds also decreased A549-ACE2 viability at micromolar concentrations similar to those where they exert their bilayer-modifying effects (**Supplementary** Figure 1C). This is consistent with previous observations showing a correlation between membrane bilayer-modifying properties and cytotoxicity.^21^

The effect of IVM on cellular health was also examined using publicly available datasets. IVM decreased cell counts after 72 h compound treatment in the Broad PRISM viability screen, a multiplexed cell proliferation assay using 480 different cancer cell lines (**Figure 2C**). Similarly, IVM caused cell death and inhibited cell growth at low micromolar concentrations after 48 h in the NCI60 tumor panel using 60 different cell lines tested in parallel (logLC_50_ = –4.5 ± 0.3, logGI_50_ = – 5.6 ± 0.1; **Figure 2D**). In neither dataset were these effects lineage-dependent. Furthermore, the IVM analogs eprinomectin and selamectin decreased cell counts in the Broad PRISM assay (**Supplementary** Figure 2A), and the IVM analogs abamectin, doramectin, emamectin, and selamectin inhibited cell growth in the NCI60 tumor panel at low micromolar concentrations (**Supplementary** Figure 2B). These previous results with IVM analogs are consistent with the cell viability counter-screens in A549-ACE cells (**Figure 1B**, **Figure 2**). The consistent adverse effects on cellular health by IVM in multiple cell lines suggest these findings are likely applicable to other cell lines including those used in previous reports with IVM in antiviral assays (e.g., Vero cells). The similar findings with IVM analogs also suggest that the observed adverse effects are common to the IVM chemotype.

The composition of complete culture media on A549-ACE2 cell viability in the presence of IVM was also evaluated. Notably, most live virus assays use 2% fetal bovine serum (FBS) to enhance viral infection, whereas most cell lines are continually propagated using 5-10% FBS. The effect of IVM on cell viability was dependent on the media FBS concentration, with nearly a log-fold difference in cell viability between media containing 0% FBS and 20% FBS even after only 24 h IVM treatment (CC_50, 24 h_ = 3.3 and 35 μM, respectively; **Supplementary** Figure 3A). This effect was observed with neither remdesivir nor staurosporine (**Supplementary** Figure 3A). It is known that IVM binds avidly to plasma proteins.^22^ In contrast to FBS, additional bovine serum albumin (BSA; 0 to 1 mM final BSA concentration) in culture media containing 2% FBS (v/v; estimated 7.5 μM final albumin concentration) did not have appreciable effects on A549-ACE2 cell viability for IVM, remdesivir, and staurosporine (**Supplementary** Figure 3B). These data suggest that BSA does not function as an additional sequestering agent for IVM. These data show that the composition of complete culture media, specifically the FBS concentration, can have a significant effect on the cytotoxicity of IVM and should be considered when comparing and interpreting cellular health and antiviral effects, and is illustrative of the more widely appreciated impact of cell culture conditions phenotypic outputs.^23^ While not investigated further, these results could be explained by growth-dependent effects (i.e., cells proliferate slower at lower FBS concentrations) and/or free-drug scavenging by lipids and proteins contained within the added FBS.

Together, these data demonstrate that IVM decreases cellular viability, induces cytotoxicity, and increases biochemical markers of apoptosis at the low micromolar concentrations, and at time points, used in SARS-CoV-2 drug repurposing assays.

### Assay interference counter-screens

A significant issue in drug discovery are nuisance compounds which can produce artifactual bioactivity by interference with assay technologies and/or true bioactivity by undesirable yet poorly translatable mechanisms of action.^24^ To gain mechanistic understanding of the *in vitro* IVM antiviral activity, we therefore evaluated IVM using a panel of assay interference counter-screens.

The original study reporting ivermectin as an NLS-importin inhibitor used the AlphaScreen homogenous proximity assay technology against a small subset of the LOPAC collection (∼400 compounds total) at a single compound concentration (10 μM).^7^ AlphaScreen technology generates an assay signal by laser excitation of a donor bead to generate singlet oxygen, which can subsequently diffuse to a proximal acceptor bead to undergo a further series of chemical reactions that results in light emission at a lower wavelength (**Figure 3A**).^25^ While reviewing the results of NCATS COVID-19 OpenData portal, we observed that IVM and analogs were active in both an AlphaLISA qHTS for inhibitors of the ACE2-Spike protein (unrelated to the purported NLS-importin target) and an AlphaLISA TruHits interference counter-screen (**Figure 3B**).^26^ Notably, these assays did not have detergent in the assay buffer, which in Alpha-based assays, can attenuate signal quenching effects by poorly soluble compounds that form aggregates.

**Figure 3.**
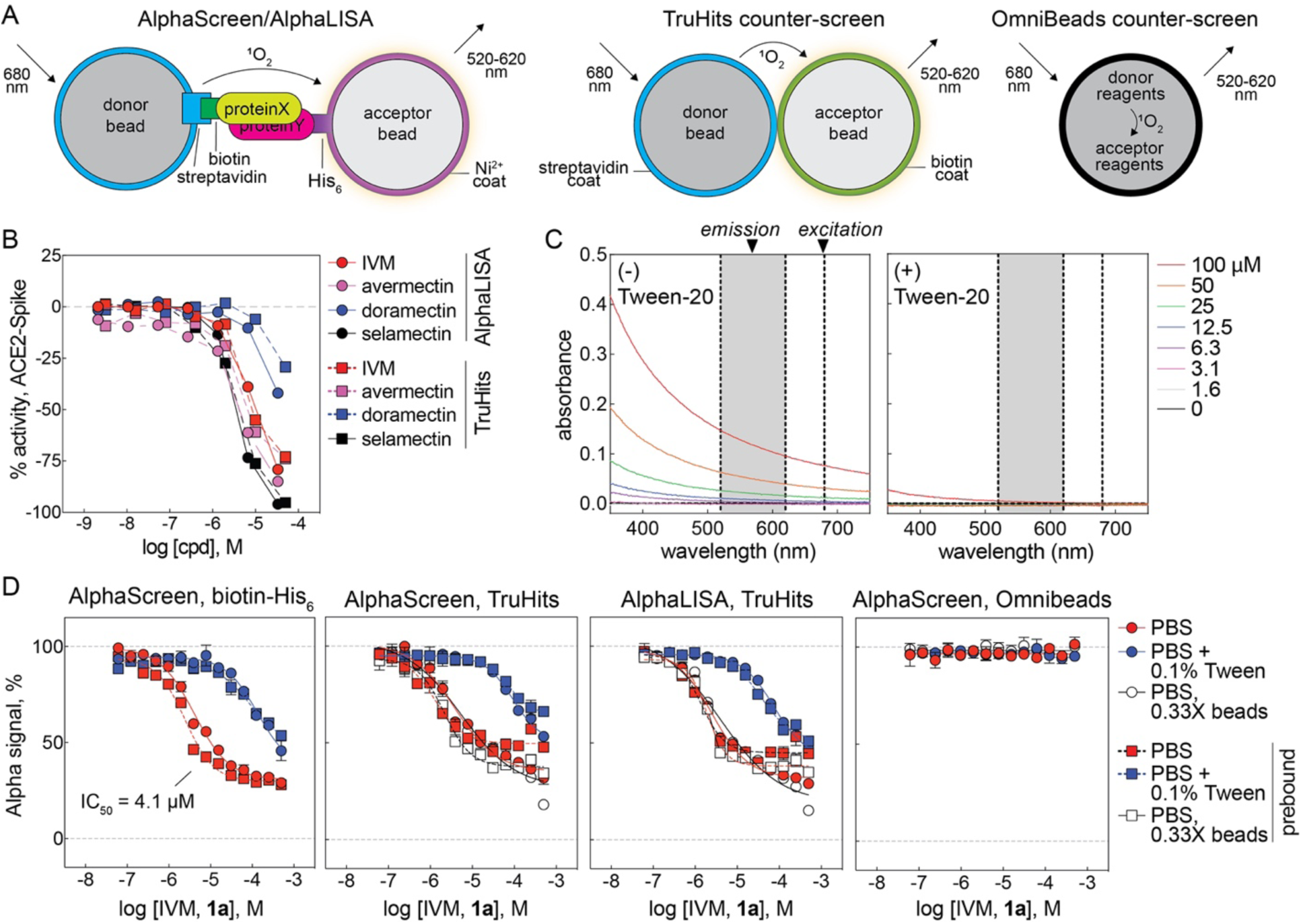
Ivermectin interferes with Alpha homogenous proximity technology by signal quenching. (A) Schematic of Alpha technology and interference counter-screens. Note that singlet oxygen (^1^O_2_) diffuses from an acceptor bead to donor bead (and can physically interact with test compound) in the AlphaScreen/LISA and TruHits formats, but not the OmniBeads format. (B) IVM and analogs decrease assay signal in an AlphaLISA qHTS for inhibitors of the ACE2-Spike interaction and a follow-up TruHits counter-screen for compound-mediated interference. Data is from NCATS COVID-19 OpenData portal. (C) UV-Vis absorbance spectrum of IVM in PBS. Data are mean ± SD of three intra-plate technical replicates. (D) IVM decreases the AlphaScreen signal in a biotin-His6 counter-screen, and the AlphaScreen and AlphaLISA signal in TruHits-based counter-screens. IVM does not affect the AlphaScreen signal in an OmniBeads counter-screen. Note the partial efficacy, which is likely due to limited solubility. Data are mean ± SD of four intra-plate technical replicates. Source data are provided as a Source Data file.

A series of counter-screens therefore were performed to characterize the potential for IVM to interfere with the AlphaScreen technology.^27^ At micromolar concentrations, the UV-Vis absorbance of ivermectin in PBS was inversely proportional to wavelength, a pattern consistent with limited solubility that results in light scattering (**Figure 3C**). Substituting the importin-NLS target with a biotinylated-His6 tag in a counter-screen designed to mimic the original AlphaScreen conditions that identified IVM as an apparent inhibitor of the NLS-importin complex effectively reproduced the original concentration-response curve (CRC) of ivermectin (**Figure 3D**). Notably, the potency and efficacy are consistent with the original AlphaScreen results for IVM versus NLS-importin (IC_50_ = 4.1 and 4.8 μM, respectively; approximately 70% maximal inhibition for both). A counter-screen similar to the one above was reported^7^, but it appears to have been performed at a single concentration, which may explain the apparent lack of compound-mediated assay interference. The addition of detergent (0.1% Tween-20 v/v) reduced the interference potency by nearly 30-fold (IC_50_ = 4.1 and 130 μM, respectively; **Figure 3D**). The addition of compound before or after the formation of the rapidly formed and high affinity biotin-streptavidin complex can help determine if a compound is interfering with the biotin-streptavidin capture reagent. In this counter-screen configuration, the interference was independent of the order of compound addition relative to the formation of the donor bead-analyte-acceptor bead complex (**Figure 3D**). Notably, IVM samples were inactive in Alpha-based PubChem screening assays containing Triton or Tween detergents (**Supplementary Table 2**). These observations are most consistent with interference in which compound aggregates physically quench singlet oxygen during its transmission from donor to acceptor bead.

IVM also reduced the signal readout in a counter-screen utilizing TruHits control beads in both the AlphaScreen and AlphaLISA formats (**Figure 3D**). These donor and acceptor beads are coated with streptavidin and biotin, respectively, and form a complex capable of generating signal without the need for an analyte to induce donor-acceptor bead proximity. As with the biotin-His6 counter-screen, the inclusion of 0.1% Tween-20 attenuated the AlphaScreen interference in the TruHits format nearly 25-fold (IC_50_ = 3.8 and 93 μM, respectively), and the interference pattern was not significantly affected by dilution of bead concentration (**Figure 3D**). These observations are also consistent with a quenching interference in the AlphaLISA technology (IC_50_ = 2.5 μM; **Figure 3D**). IVM did not interfere with the assay signal in a counter-screen utilizing OmniBeads, a single reagent bead that contains all of the Alpha chemistry reagents and therefore does not require two proximal complementary beads to produce an assay signal (**Figure 3D**). This observation is again consistent with IVM interference by quenching the donor-to-acceptor signal transmission, as the test compound is not able to quench the intra-bead singlet oxygen.

The control compounds biotin (capture reagent mimetic), GW-5074 (light scatterer), piceatannol (singlet oxygen quencher), and methylene blue (colored) in these counter-screens all behaved consistent with their expected modes of AlphaScreen technology interference (**Supplementary** Figure 4)^27^. Notably, another compound identified in the same report as ivermectin, mifepristone (which is also poorly soluble in water^28^), also interfered with the AlphaScreen technology readout in a manner consistent with signal quenching (**Supplementary** Figure 4). Together, these results demonstrate that IVM interferes with the AlphaScreen homogenous proximity assay technology used in its original repurposing.

IVM was profiled in several counter-screens for colloidal aggregation, a common compound-mediated assay interference in biochemical and occasionally cellular assays. Presumably due to its high ClogP, IVM was flagged as a potential aggregator in two predictive models: Aggregator Advisor and SCAM Detective (**Supplementary** Figure 5A and 5B).^29, 30^ However, in the absence of nonionic detergent, IVM did not inhibit AmpC β-lactamase or malate dehydrogenase activity at micromolar concentrations, two unrelated enzymes sensitive to aggregators (**Supplementary** Figure 5C). In a previous qHTS without detergent, IVM was an apparent activator of cruzain, another prototypical enzyme sensitive to aggregators (AC_50_ 4.98 μM, efficacy 165%; PubChem AID 1476, CID 11957587). Using dynamic light scattering, IVM only formed colloidal aggregates at high micromolar concentrations (approximately 100 μM) well above its solubility limit of 1-2 μM (**Supplementary** Figure 5D).^10, 11^ As a whole, these data suggest small-molecule colloidal aggregates, which can nonspecifically perturb protein function, are unlikely to be responsible for the IVM antiviral and cytotoxic activities at low micromolar concentrations.

IVM was also profiled in a counter-screen for drug-induced phospholipidosis (DIPL), another common compound-mediated assay interference in cellular assays usually associated with cationic amphiphiles. IVM did not induce detectable levels of DIPL in A549-ACE2 cells, and only induced DIPL at high micromolar concentrations in Hep G2 cells (**Supplementary** Figure 5E). These data suggest DIPL is unlikely to be primarily responsible for the IVM antiviral and cytotoxic activities typically observed at low micromolar concentrations. This is not unexpected, as IVM does not contain the prototypical cationic amphiphilic moieties, like protonated amines, that are associated with many DIPL compounds.^31^

The effect of IVM on several SARS-CoV-2-related molecular targets was also examined using data from the NCATS COVID-19 OpenData portal.^26^ IVM did not inhibit human ACE2 enzymatic activity, SARS-CoV-2 3CL activity, human TMPRSS2 activity, or SARS-CoV-2 RdRp activity in biochemical assays, even at 50 μM concentrations (**Supplementary** Figure 6).^26^ Notably, none of these assays involved membrane-based targets.

### Off-target profiling

The classical vermicidal mechanism of IVM (at low nanomolar concentrations) is due to IVM binding to chloride-conducting glutamate-gated Cys-loop receptors, Glu-Cl,^32^ which activates the channels and paralyzes the worms. The apparent antiviral effects and cellular injury produced by IVM, in contrast, occur at micromolar concentrations. Given that IVM is known to alter the function of many different membrane proteins at micromolar concentrations (**Supplementary Table 1**), we hypothesized that the classical mechanism of action for IVM (at nanomolar concentrations) could be a clue as to the molecular mechanism of its apparent antiviral effects and accompanying cellular injury (at micromolar concentrations). It is in this context likely to be relevant that IVM is a hydrophobic macrocyclic lactone (ALogP estimates vary between 4.4 and 5.8; the experimental octanol/water partition coefficient is ∼1,600)^33, 34^ that partitions into the outer leaflet of cellular membranes^6^, where it can bind to membrane-based targets (**Figure 4A**). This raises the possibility that IVM at micromolar concentrations may accumulate sufficiently in membrane bilayers to perturb the function of embedded proteins by a bilayer-mediated regulation, and perhaps engage membrane-based targets by low-affinity interactions, similar to what has been reported for platelet-activating factor.^35, 36^

**Figure 4.**
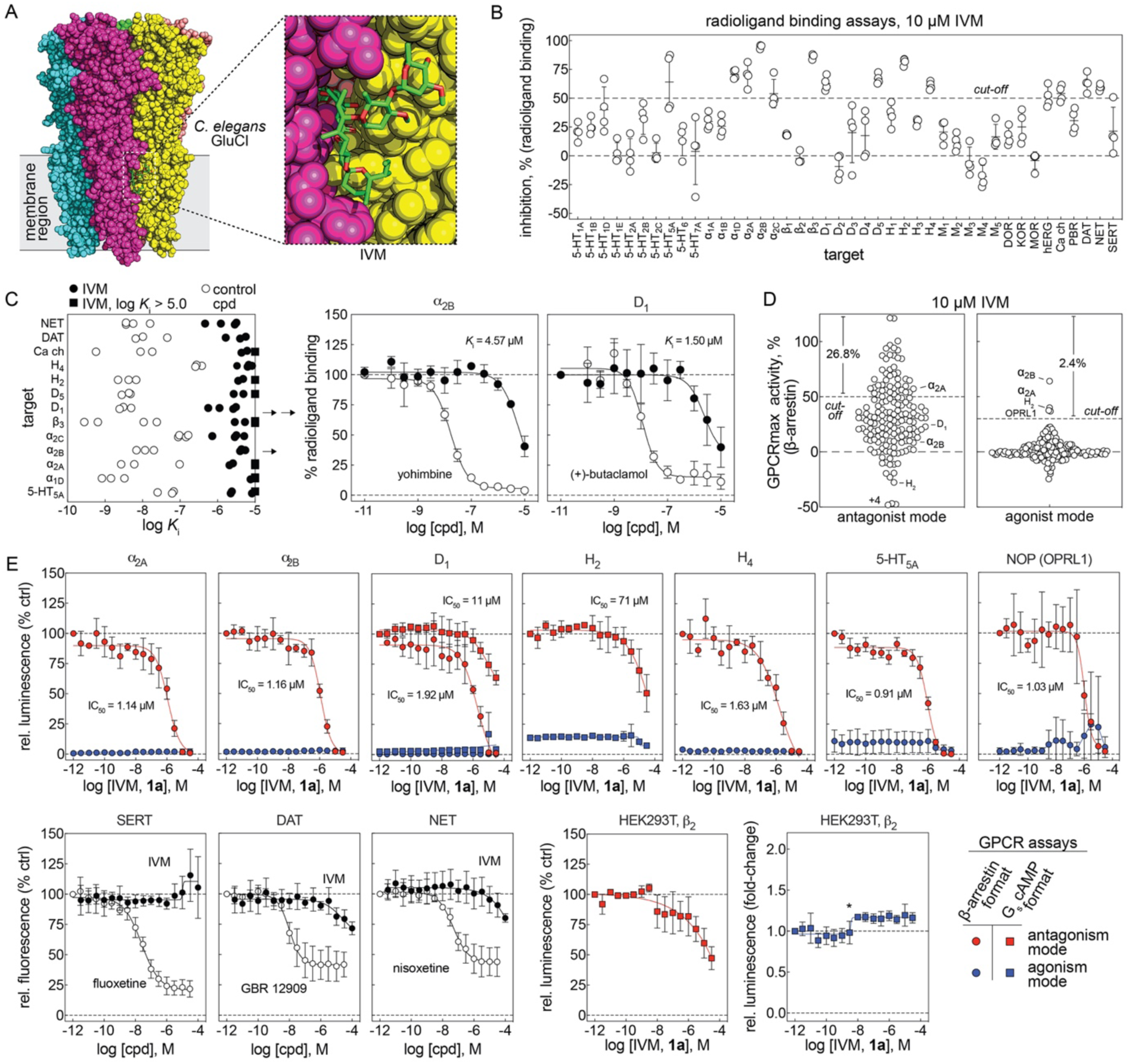
Ivermectin nonspecifically perturbs membrane-based targets including GPCRs. (A) IVM antiparasitic activity involves compound binding at the membrane-membrane protein interfaces (e.g., *C. elegans* glutamate-gated chloride channel, PDB 3RHW)^32^. (B) IVM nonspecifically inhibits radioligand binding to membrane-bound targets including GPCRs. Data are four intra-plate technical replicates with errors bars denoting mean ± SD. (C) Calculated log K_i_ values of IVM from confirmatory radioligand binding concentration-response testing with each point representing an independent experiment. Reference control compounds were included in each experiment. Inset: select concentration-response curves; data are mean ± SD of three independent experiments. (D) IVM nonspecifically antagonizes GPCR function in a cellular β-arrestin reporter assay (gpcrMAX panel, Eurofins). Data are mean of two technical replicates. (E) IVM nonspecifically antagonizes GPCR function in follow-up concentration-response testing in cellular functional assays. IVM was tested in both agonist and antagonism modes in β-arrestin (Tango) and/or G_s_ cAMP (GloSensor) formats. IVM was also tested for inhibition of membrane-based transporters. Data are mean ± SD from three to four independent experiments. Details of reference agonists and their concentrations are provided in Source Data. *, confirmed detector-based artifact at low luminescence (baseline HEK293T antagonism = ∼3000 RLU, baseline HEK293T agonism = ∼50 RLU). Source data are provided as a Source Data file.

Because IVM perturbs membrane bilayers at concentrations similar to those where it inhibits viral replication and cellular health, it was tested for its ability to perturb membrane-based targets. Given its association with neurological adverse events^37^, IVM was profiled by the Psychoactive Drug Screening Program (PDSP) for binding to a panel of primarily G-protein-coupled receptors (GPCRs) by radioactive ligand-binding assays using membrane-bound GPCRs, in the form of crude membrane preparations from transiently transfected or stable cell lines (HEK293 or CHO).^38^ Using a cut-off of 50% inhibition, IVM significantly decreased radioligand binding in 14/42 (33%) of the targets profiled at 10 μM (**Figure 4B**). Follow-up concentration-response assays demonstrated that IVM inhibited these targets with affinities (*K*_i_ values) in the same low micromolar ranges where it displays apparent antiviral activity and produces cellular injury (**Figure 4C**). Notably, the assay readout does not inform *how* the compound inhibits ligand binding to these targets (direct competition with the radiolabeled ligand or a more nonspecific inhibitory mechanism).

IVM was also profiled at 10 μM for functional agonist and antagonist activity in a panel of 168 GPCRs using a cellular β-arrestin reporter assay.^39^ Consistent with the (nonspecific) activity observed in the PDSP profiling, IVM inhibited the β-arrestin readout in 26.8% (45/168, 50% antagonism cut-off) of the tested GPCRs (**Figure 4D**). By contrast, IVM activated the β-arrestin readout in only 2.4% (4/168, 30% agonism cut-off) of the tested GPCRs (**Figure 4D**). The sheer number of GPCRs antagonized by IVM is consistent with a nonspecific mechanism of action in cells.

For select targets identified through either the binding or functional profiling studies, the agonist and antagonism activity was further assessed with secondary (concentration-response) functional assays: a β-arrestin reporter (Tango assay) and a cAMP biosensor (GloSensor assay).^40, 41^ IVM antagonized the function of several GPCRs at low micromolar concentrations in the β-arrestin reporter format, whereas the antagonism by IVM appears less potent in the cAMP reporter format (**Figure 4E**). Notably, the α_2B_ receptor showed both antagonism and agonism in the primary GPCR functional screen, but follow-up testing was consistent with receptor antagonism. Furthermore, IVM displayed inhibitory activity in a control assay in parental HEK293T cells when endogenous β_2_ receptors were stimulated by isoproterenol and measured using the cAMP biosensor, suggesting an inhibitory activity at β_2_ receptors. By contrast, treatment with IVM did not lead to significant stimulation of cAMP production in HEK293T cells (**Figure 4E**). It is unlikely that IVM interfered with the luciferase-based reporters, as IVM and a close analog avermectin A_1a_ did not inhibit several recombinant luciferases in previous qHTS experiments (**Supplementary** Figure 7). Different from its effect on GPCRs, IVM had little inhibitory activity on several tested neurotransmitter transporters (DAT, NET, and SERT; **Figure 3E**). Consistent with previous studies (**Supplementary Table 1**), these results suggest that IVM can promiscuously inhibit membrane-based protein targets, including GPCRs, at the same concentrations where it exerts antiviral activity and causes cellular injury.

### Analysis of PubChem data

The activity of IVM samples tested in PubChem screening assays (n = 766) was evaluated for bioassay promiscuity and trends related to assay design. Remarkably, nearly all of the PubChem screening assays in which IVM was classified as active were cellular assays and not biochemical assays (**Table 1**). Many of the cellular assays explicitly assayed either cell growth, cell viability, or cytotoxicity (153/493, 31.0%), and both the toxicity-related and the non-toxicity related cellular assays had high activity rates (31.4% and 17.9%, respectively). The activity classifications between biochemical and cellular assays were not evenly distributed, even when toxicity-related cellular assays were excluded (both analyses p < .0001). The high activity rate in all PubChem cellular assays (104/493, 22.1%) compared to biochemical assays (2/273, 0.73%) also suggests that IVM is promiscuous in high-throughput cellular assays performed with low micromolar concentrations of IVM. These observations further suggest that membrane bilayers are a key component for IVM bioactivity.

**Table 1.** Summary of ivermectin bioactivity in PubChem. Assays in which IVM is classified as active are enriched in PubChem cellular screening assays compared to biochemical assays. The activity classifications between biochemical and cellular assays (all cellular assays and non-toxicity related assays) were not evenly distributed (null hypothesis rejected). Source data provided as a Source Data file.

| | | PubChem activity annotation | | | $\chi^2$ (compared to biochemical assays) |
| --- | --- | --- | --- | --- | --- |
|  |  | active<sup>1</sup> | inactive | other<sup>2</sup> |  |
| assay category | biochemical | 2 | 246 | 25 | - |
| | cellular (all) | 109 | 322 | 62 | $\chi^2(2, N = 766) = 71.78, p < .0001$ |
| | cellular (non-toxicity related subset) | 61 | 229 | 50 | $\chi^2(2, N = 613) = 57.56, p < .0001$ |

<sup>1</sup>, active or probe; <sup>2</sup>, inconclusive or unspecified

### Membrane bilayer perturbation

To further explore the possibility that IVM may produce some of its effects through bilayer-mediated regulation of membrane protein function, IVM was tested for membrane bilayer perturbation using a gramicidin-based fluorescence quench (GFA) assay. The assay takes advantage of thallium ion (Tl^+^) permeability through bilayer-spanning gramicidin channels, to quench the fluorescence of the 8-aminonaphthalene-1,3,6-trisulfonic acid (ANTS) fluorophore. Gramicidin channels form by transmembrane dimerization of non-conducting subunits^42^, which are incorporated into the membrane of ANTS-loaded large unilamellar vesicles (LUVs). When the channels’ hydrophobic length (∼2.3 nm)^43, 44^ is less than the thickness of their host bilayer (∼3.4 nm^45^, in the case of the 1,2-dierucoyl-sn-glycero-3-phosphocholine membranes used in this study) channel formation produces a local bilayer deformation/thinning. This deformation incurs an energetic cost, which becomes the bilayer contribution 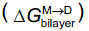 to the total free energy of dimerization 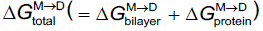, where the superscripts “M” and “D” denote the gramicidin monomers and dimers, respectively, and the subscripts “total” and “protein” denote the total energy and the energetic contribution from rearrangements within the gramicidins as they dimerize. Given the gramicidin channels’ structure, being near-cylindrical, drugs are unlikely to bind to the bilayer-spanning dimer.^46, 47^ When amphiphiles partition into the bilayer/solution interface, they will alter lipid bilayer properties (thickness, intrinsic curvature and the associated elastic moduli)^48^, which will alter (usually decrease) 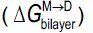 and in turn the gramicidin monomerΗdimer equilibrium (usually toward the right). The shifts in the gramicidin monomerΗdimer equilibrium can be measured from the drug-induced shifts in the rate at which extravesicular Tl^+^ enters LUVs through gramicidin channels and quenches the ANTS fluorescence (quantified as the relative changes in rate, *rate*/*rate*_control_). Drug-induced changes in the ANTS quench rate thus reflect the bilayer-modifying potency of a compound,^49, 50^ and changes in *rate*/*rate*_control_ > 1.5 are associated with increased risk that the compound may be cytotoxic.^21^ In initial testing, IVM significantly increased the ANTS quench rate at 10 µM (**Figure 5A**). In concentration-response testing, IVM increased the ANTS quench rate in a concentration-dependent manner, with 1 μM IVM increasing the rate relative to DMSO control 12-fold indicating that IVM is a potent bilayer modifier at this concentration (**Figure 5B**). For comparison, only 1/400 (0.25%) drugs in Medicines for Malaria Venture’s Pathogen Box increased the quench rate more than 10-fold at a higher 10 µM concentration, and only 33/400 (8.2%) drugs increased the quench rate more than three-fold at 10 µM.^21^ IVM is an exceptionally potent bilayer-modifying molecule. In contrast, remdesivir had no observable effect at 1 µM, and only caused modest two-fold changes in quench rate at 3 μM (two-fold the IC_50_ for inhibition of viral growth in the SARS-CoV-2 FLuc assay; **Figure 5B**). Remdesivir thus would be expected to be (much) less cytotoxic than IVM, as observed (**Figure 1A**). We also tested a blinded (to the experimenter) library of IVM analogs and remdesivir at 10 µM concentration (**Supplementary Table 3**); the results are unmarkable (flat SAR), except for the avermectin B_1a_ aglycone (**12**), which has a two-fold lower bilayer-modifying potency than avermectin B_1b_ **10** (but the ivermectin B_1a_ aglycone **3** is equipotent to IVM and the ivermectin B_1a_ monosaccharide **4**).

**Figure 5.**
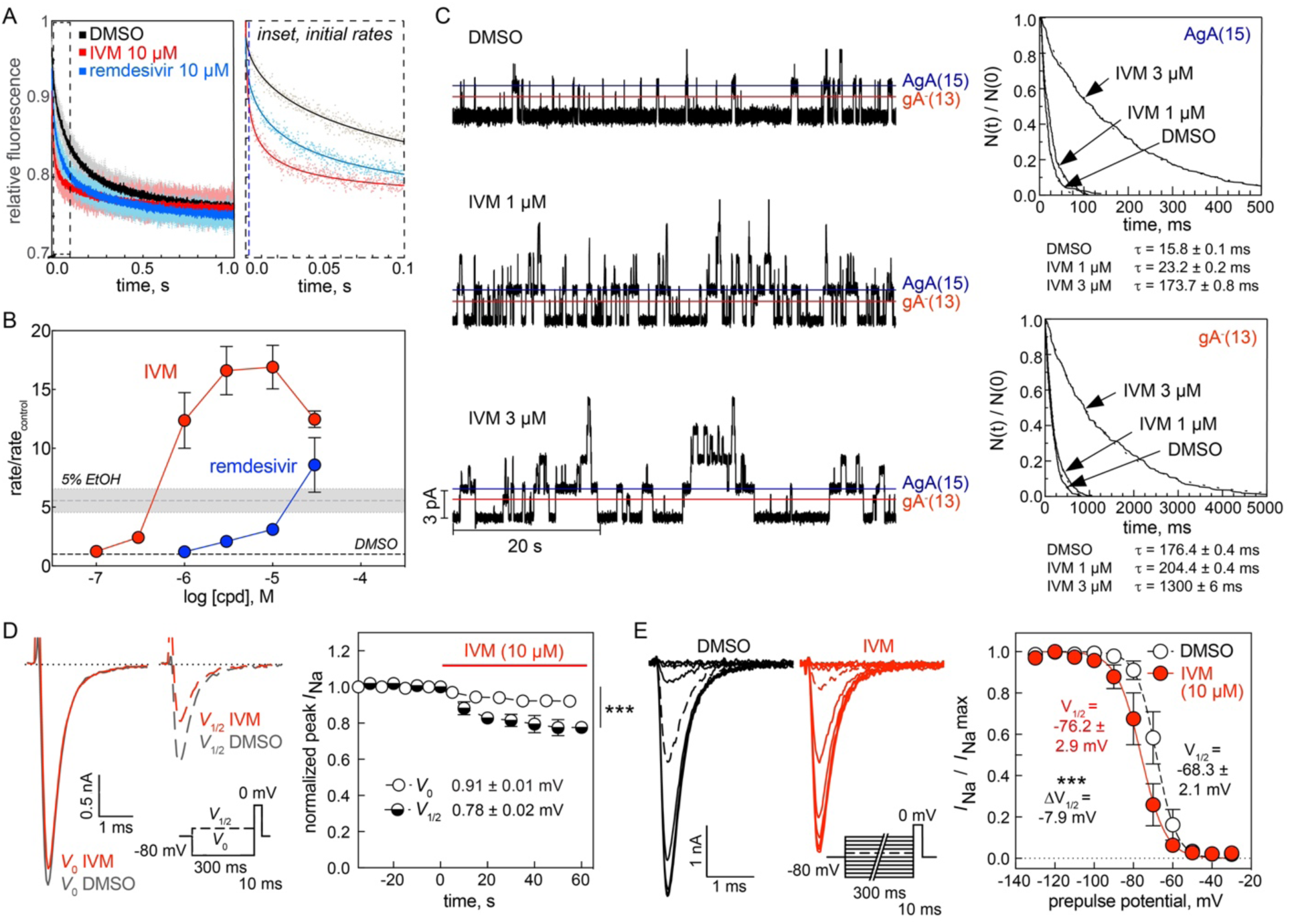
Ivermectin nonspecifically perturbs lipid bilayers and ion channels. (A) A gramicidin-based fluorescence assay (GFA) demonstrating how the influx of the heavy ion quencher Tl^+^ into fluorophore-loaded LUVs increases with IVM (**1a**) or remdesivir. Left: stopped-flow fluorescence quench traces of normalized fluorescence decay over 1 s with light dots representing individual replicates (n = 9 technical replicates) and dark dots representing replicate mean. Right: representative time courses of the initial 100 ms with dots denoting individual time points and solid lines denoting exponential fits (2–100 ms). (B) GFA concentration-response curves demonstrate IVM significantly perturbs membrane bilayers at low micromolar concentrations. 5% EtOH, positive control. Data are mean ± SD for three biological replicates (independent LUV batches). (C) IVM induces changes in gramicidin A (gA) function. Left: representative single-channel traces recorded in a planar bilayer doped with the right-handed AgA(15) and the left-handed gA^−^(13) analogs, which form channels of different lengths and helix sense. Note that 3 µM IVM produces increases in channel lifetimes and a decrease in channel appearance rate. Right: representative single-channel survivor plots for gA^−^(13) channels (top) and AgA(15) channels (bottom) with single-exponential fits to the survivor distributions. IVM produced concentration-dependent increases in the average single-channel lifetimes (1). Data are from one of seven independent experiments each with at least three separate measurements. See also **Table 2** for experimental summary. (D) IVM causes voltage-dependent inhibition of the peak Na^+^ current. Time-course of drug wash-in showing inhibition of voltage-dependent sodium channels in neuroblastoma ND7/23 cells using an alternating two-pulse protocol. Left: representative Na^+^ current traces from an alternating two-pulse protocol. A test pulse to 0 mV to elicit peak Na^+^ current (*I*_Na_) was preceded by a 300 ms prepulse to holding potentials of either V_0_ (−130 mV, solid lines) or V_½_ (dashed lines, see insert; V_½_ = −69 ± 3 mV). Black lines, DMSO control; red lines, 10 µM IVM. IVM significantly reduced peak I_Na_ to 0.91 ± 0.01 mV (prepulse to V_0_, *p* < 0.0001) and peak I_Na_ to 0.78 ± 0.02 mV (prepulse to V_½_, *p* < 0.0001). Data are mean ± SD from four to six biological replicates. (E) IVM promotes inactivation of voltage-dependent sodium channels. Left: representative Na^+^ current traces from a double-pulse protocol in which a test pulse to 0 mV was preceded by a 300 ms conditioning prepulse to potentials ranging from –130 mV to –30 mV. The data from each experiment were fitted with a standard two-state Boltzmann equation to calculate the individual V_½_ values. Right: fitted data showing IVM causes a hyperpolarizing shift of the voltage-dependance of half-maximal inactivation (ΛV_½_ = –7.9 mV, *p* < 0.0001). Data are mean ± SD from four to six biological replicates. Source data provided as a Source Data file.

**Table 2.** Summary of ivermectin-induced changes in gramicidin channel function. Data are weighted mean ± SD from three to seven biological replicates (days), with the weight function being the number of events (current transitions or lifetimes) recorded in three to seven individual measurements in that experiment, which ranged between 242 and 2991 for the conductance measurements and 141 and 1651 for the lifetime measurements. See also **Figure 5C**. Source data provided as a Source Data file.

| IVM concentration (μM) | single-channel conductance (pS) | | single-channel lifetime (ms) | | lifetime ratio ($\tau$) | normalized lifetime ratio |
| --- | --- | --- | --- | --- | --- | --- |
| | Gly<sup>-</sup>(13) | AgA(15)15 | Gly<sup>-</sup>(13) | AgA(15)15 | ($\tau_{13}/\tau_{15}$) | $\frac{(\tau_{13}/\tau_{15})_{drug}}{(\tau_{13}/\tau_{15})_{ctrl}}$ |
| 0 (DMSO) | 11.1 ± 1.0 | 17.5 ± 1.2 | 15 ± 2 | 167 ± 11 | 0.09 ± 0.02 | 1.00 |
| 1 | 10.4 ± 1.0 | 16.9 ± 1.3 | 55 ± 19 | 350 ± 134 | 0.14 ± 0.03 | 1.41 ± 0.23 |
| 3 | 10.5 ± 0.8 | 16.7 ± 0.9 | 139 ± 44 | 973 ± 425 | 0.15 ± 0.03 | 1.67 ± 0.61* |

\*: $p = 0.049$ relative to 0 μM IVM (DMSO control) using a two-tailed Mann-Whitney test

In addition to the fluorescence quench-based assay, we examined the action of IVM using single-channel electrophysiology, which provides for additional information about bilayer-perturbing effects.^51, 52^ Using planar bilayers formed from a suspension of 1,2-dioleoyl-*sn*-glycero-3-phosphocholine in *n*-decane, we examined the effects of IVM on gramicidin channels formed by the 15-amino-acid right-handed [Ala^1^]gramicidin A (AgA(15)) and the chain-shortened, left-handed des-Val-Gly-gramicidin A with 13-amino-acid (gA^−^(13)). These gramicidin analogues form right– and left-handed channels, respectively, that differ in length by 0.32 nm, and the bilayer deformation (and bilayer deformation energy) associated with the shorter gA^−^(13) channels will be larger than that associated with the longer AgA(15) channels. The bilayer responds to the deformation by imposing a force on the bilayer-spanning channel, which will promote the dissociation of the bilayer-spanning channel, with the greater force acting on the shorter channel. The opposite handedness prevents heterodimerization, which simplifies the analysis, and the homodimeric channels can be distinguished by their characteristic single-channel current transition amplitudes, as shown in single-channel current traces (**Figure 5C**, left). As summarized in **Table 2**, 1 µM IVM increased the durations of the conducting events for both the left– and the right-handed channels with little effect on the single-channel conductances, which indicates that direct interactions with the gramicidin channels are unlikely. At 3 µM, however, IVM produced further increases in the average channel lifetimes, but decreased the appearance frequencies and began to destabilize the membranes. Importantly, the relative increases in the average channel lifetimes (τ) were larger for the shorter gA^−^(13) channels, as compared to the longer (AgA(15) channels (**Figure 5C**, right). These observations with IVM in gramicidin A are consistent with previously characterized amphiphiles that increase membrane bilayer elasticity at low micromolar concentrations.^52, 53^

A general feature of molecules that increase bilayer elasticity is that they promote the inactivation of voltage-dependent cation channels.^52, 54, 55, 56^ We therefore tested whether IVM promotes the inactivation of voltage-dependent sodium channels (Na_V_) as another surrogate for membrane perturbation. The distribution between activatable and inactivated Na_V_’s varies with membrane potential, with depolarization promoting inactivation. We used an alternating pulse protocol (**Figure 5D**, inset) to test for voltage-dependent effects after a prepulse to either –130 or to *V*_½_ (the potential where approximately half the channels were in the inactivated state; –69 ± 3 mV, a separate curve was obtained for each cell tested).^56, 57^ As expected, IVM produced voltage-dependent inhibition of the peak Na^+^ current (*I*_Na_) to (**Figure 5D**). IVM decreased the peak *I*_Na_ to 0.91 ± 0.01 (prepulse to V_0_, *p* < 0.0001). As is the case for other amphiphiles, the inhibition was greater when the test pulse was preceded by a prepulse to *V*_½_ (the potential where approximately half the channels were in the inactivated state).^52, 57^ In this case, IVM decreased the peak *I*_Na_ to 0.78 ± 0.02 (prepulse to *V*_½_, *p* < 0.0001).

The effect of IVM on Na_V_ steady-state inactivation (often referred to as Na^+^ channel availability) was also explored using a double-pulse protocol (inset in **Figure 5E**). This protocol increases the number of channels in the inactivated state with each successive pulse, which tests the ability of the IVM to stabilize the inactivated state which will be reflected in a shift of the steady state curve toward more hyperpolarized potentials. IVM does indeed promote Na_V_ inactivation (**Figure 5E**), as would be expected for a compound that increases bilayer elasticity. As is the case for other compounds that increase bilayer elasticity^52, 56, 57^, IVM does indeed promote a hyperpolarizing shift in the Na_V_ availability curve (βV_½_ = −7.9 mV, *p* < 0.0001). Compared to other amphiphiles like phytochemicals or volatile anesthetics, however, the shift produced by IVM is modest; amphiphiles that produce a three-fold change in quench rate tend to produce a shift in *V*_½_ by about 10-20 mV, whereas 10 µM IVM produced a 17-fold increase in quench rate and a −7.9 mV shift in *V*_½_. The seeming lesser potency of IVM on Na_V_ availability may reflect that it resides in the outer leaflet of the plasma membrane^6, 13^, whereas the molecule motions that underlie voltage-dependent gating seem to occur in the inner leaflet.^58, 59^ Together, the results from both the alternating pulse and steady-state inactivation protocols demonstrate that IVM disrupts the function of voltage-gated sodium channels, and does so in a manner consistent with other previously characterized small-molecules that perturb membrane bilayers.

## Discussion

Given the reported interest in the antiparasitic drug IVM in the treatment of SARS-CoV-2, we evaluated IVM for its potential to be repurposed as a SARS-CoV-2 agent using several preclinical assays. Consistent with the original report^8^, IVM was able to reduce SAR-CoV-2 virus levels in human cells at low micromolar concentrations. This antiviral activity correlated with multiple markers of cellular injury including decreased cell viability, cytotoxicity, and apoptosis. Drug-induced phospholipidosis and colloidal aggregation, two common mechanisms of cellular assay interference, were ruled out as causative mechanisms for the cellular injury. However, IVM was found to perturb membrane bilayers, promote Na_V_ inactivation, and nonspecifically engage membrane-based targets such as GPCRs and noncanonical ion channels at low micromolar concentrations. Moreover, IVM was found to grossly interfere with AlphaScreen homogenous proximity assays, the screening technology originally used to identify IVM as a purported NLS-importin disruptor.

The effects of micromolar concentrations of IVM on cells are consistent with previous reports of promiscuous membrane bilayer effects associated with many drugs and phytochemicals.^21, 36, 60^ Some bilayer-perturbing compounds may do so through dramatic bilayer destabilizing/reordering mechanisms especially at high concentrations.^61^ We focus here on more subtle bilayer effects, which do not disrupt the bilayer but alter its elasticity, curvature and/or thickness.^62, 63^ Using fluorescence quench experiments we show that IVM shifts the gramicidin monomerΗdimer equilibrium toward the conducting dimers. Using single-channel electrophysiology we show that IVM causes larger relative changes in the average lifetimes of the shorter gA^−^(13) channels than of the longer VgA(15) channels, which shows that IVM increases the bilayer elasticity.^53, 64^ Off-target and/or toxic (pleiotropic) actions of certain drugs have been found to occur at bilayer perturbing concentrations.^36, 52, 64^ Moreover, such compounds tend to alter the function of multiple membrane proteins at bilayer-perturbing concentrations, which is evident in the GPCR profiling of IVM (**Figure 4**). We show that IVM, like many other amphiphiles, promotes Na_V_ inactivation, with greater potency when function is tested from more depolarized potentials (**Figure 5E**). The shift in the inactivation curve relative to the IVM-induced change in *Rate*/*Rate*_control_ is less than that observed with other drugs, e.g., certain volatile anesthetics^57^, which may reflect that IVM is likely to reside primarily in the outer leaflet of the plasma membrane^6^ whereas the gating motions in voltage-dependent sodium channels occur toward the cytoplasmic end of the channel.^58, 59, 65^ The generality of such “membrane effects” is evident also in IVM’s agonist and antagonist effects on GPCR’s (**Figure 3**). The overlap between the concentrations at which drugs have pleiotropic effects and nonspecifically regulate structurally and functionally diverse membrane proteins and concentrations at which they alter the lipid bilayer properties suggests a common mechanism for off-target effects that is mediated by the lipid bilayer.^36^

Our results are consistent with more physiologic models of coronavirus infection, which also showed a lack of efficacy in a human-airway-derived cell model.^66^ In general, there have been mixed results in animal models assessing IVM as an anti-coronaviral agent. A study on mouse hepatitis virus (single 500 µg/kg dose of IVM) demonstrated some reduction in viral load.^67^ A study (400 µg/kg of IVM per day over four days) showed an overall lack of efficacy in Syrian hamnsters.^68^ Another study in hamsters showed no decrease in viral load but reported attenuation of clinical and immunological outcomes by invoking a potential immunomodulatory mechanism distinct from the original hypothesized importin target.^69^ Our work suggests that IVM, and other compounds with similar modes of action, could be flagged earlier in the translation process, perhaps prior to transitioning from culture monolayers to more physiological models (e.g., which incorporate apical air and basolateral medium compartments) by testing for their (subtle) membrane-perturbing effects.

Several cell-based experiments have been used to support the proposed NLS-importin mechanism of antiviral action, including imaging-based assays with fluorophore-labelled NLS proteins.^7, 9, 70^ Biophysical techniques including thermostability, analytical centrifugation, and circular dichroism have also been used to support this mechanism, including that IVM alters protein conformation.^9^ Many of these experiments involved concentrations of IVM (> 10 μM) that are much higher than the reported solubility limits^10, 11^, which likely would lead to gross cell injury— though some involved unconventionally short treatment times. Biophysical assays can also produce positive readouts at supraphysiologic concentrations, including by nonspecific protein binding. We also note that cell-based readouts can result from desirable as well as undesirable mechanisms of action (e.g., is the reduction of viral load a direct action of IVM or a consequences of poor cell health?), and are susceptible to confirmation bias when using high compound concentrations.^24^ The data presented here suggest that the cellular assay nuclear importin readouts may be confounded by pleiotropic effects due to off-target effects from IVM.

Our results are also consistent with emerging clinical data. Recent reports have shown a lack of clinical efficacy of IVM for the treatment of SARS-CoV-2 in high-quality clinical trials (**Supplementary Note 2**). There are multiple hypotheses for a lack of observed efficacy, including pharmacokinetics such as insufficient dosing or suboptimal timing in relation to disease occurrence or viral exposure. There have been proposals for alternate strategies such as IVM analogs, alternative formulations (e.g., aerosolized IVM), and alternative dosing regimens. Based on this work and the cumulative evidence to date, however, it is likely that such strategies would be ineffective and perhaps cause harm. It is possible, however, that IVM may be helpful in populations with a high incidence of helminthic diseases, where IVM in effect could decrease the incidence of comorbidities.

The use of the gramicidin assay as a surrogate for nonspecific membrane bilayer modification could become a robust frontline assay for evaluating both repurposed drugs as well as cell-active compounds identified through conventional real and virtual screening campaigns. This would be analogous to the routine counter-screening of bioactive compounds by aggregation, cytotoxicity, and reactivity counter-screens. This medium-throughput assay could be offered by contract research organizations (CROs), for example, or offered as a service by non-profit drug discovery initiatives.

This work adds to a growing list of examples of suboptimal chemical matter that has been proposed as SARS-CoV-2 therapeutics. The list includes several proposed SARS-CoV-2 main protease inhibitors that are nonspecific electrophiles^71^, cationic amphiphiles whose antiviral activity correlates with drug-induced phospholipidosis^31^, and colloidal aggregators in biochemical SARS-CoV-2 drug repurposing assays^72^. Fortunately, many of the preclinical tools and workflows used to evaluate IVM as an antiviral are public knowledge. Pertinent to the case of IVM, the NCATS *Assay Guidance Manual* contains several chapters devoted to cellular health assays, and homogenous proximity assay interferences.^73^ Unfortunately, it appears these tools are not being used to their full potential. This work underscores the need to revisit research carried out at the height of the pandemic and therefore, with a great sense of urgency but perhaps done not as rigorously as required. This is a serious issue, as evidenced by recent critiques of drug repurposing related to the pandemic and more generally.^74, 75^ As shown in the present study, as well as previous studies^31,71,72^, a key consideration in any drug repurposing effort will be to ensure that the drug is safe and efficacious at the concentrations that would need to be employed in clinical settings. It may in this context be important that the bilayer-modifying potency of a drug might be minimized without affecting its desired effects, though the flat SAR for IVM observed in the SARS-CoV-2 FLuc and CTG assays (**Figure 1D**) are not encouraging for such efforts.

There are important caveats to this preclinical evaluation. First, it remains possible that under certain experimental conditions, the antiviral effect of IVM may occur at lower concentrations than it induces cellular injury. An unforgiving dependence on conditions, however, bodes poorly for translatability. Second, it is possible that IVM may disrupt the NLS-importin function as originally proposed. However, it is again unlikely to be translatable because IVM does not appear to cross plasma membranes^6^, it is not found in organellar membranes, and any disruption of NLS-importin function would likely occur at high micromolar concentrations of IVM, which would be highly unlikely to be achieved without serious toxicity and without invoking localized intracellular accumulation.

This study provides a preclinical framework for evaluating drug repurposing candidates that ideally should be performed prior to animal studies and clinical trials, or during times of urgency such as pandemics, concurrently with existing clinical trials. In cases where the proposed repurposed activity is not reproducible, or is likely to occur by a poorly translatable mechanism, these studies could serve as a “no-go” or “halt” decision.^24^ Based on our results, the following experiments should be prioritized for any drug that is being contemplated to be repurposed: (1) reproduction of the original repurposing studies and orthogonal confirmation^76^, (2) cellular health studies at multiple concentrations and extended time points^24^, (3) counter-screens for common modes of biochemical and cellular assay interferences, (4) off-target profiling, and (5) determination of bilayer-perturbing potency. Many of these studies can be accomplished in parallel by small teams in a matter of months with sufficient funding, free access to collaborators, and outsourcing to CROs. A barrier to this model is the lack of a leading authority to guide these studies. Currently, most academic labs do not have the necessary resources or expertise to perform these studies and industry may lack financial incentives.

## Materials and methods

### Compounds and reagents

Test compounds were typically prepared as 10 or 25 mM stock solutions dissolved in neat DMSO and stored at –30 °C until use. Concentrated IVM stock solutions were prepared to obtain complete concentration-response curves. No gross precipitation of IVM was observed in DMSO up to 50 mM final concentrations, even after repeated freeze-thaw cycles. All compounds were subjected to internal quality control; most demonstrated greater than 95% purity and detection of an expected parent ion by UPLC-MS (UV, ELSD, ESI-MS; **Supplementary Table 4**). The primary IVM sample (**1a**; USP standard) was further characterized by ^1^H NMR, ^13^C NMR, UPLC-MS, and an *in vivo Caenorhabditis elegans* assay (**Supplementary** Figure 8).

### Cell lines, viruses, and cell culture

Cells were obtained from the following sources: A549-ACE2 (courtesy of Pei-Yong Shi, UTMB), A549-ACE2-TMPRSS2 (courtesy of Carol Weiss, US Food and Drug Administration), and Hep G2 (ATCC, cat # HB-8065). Cells were maintained in DMEM (Gibco, cat # 11965) supplemented with 10% fetal bovine serum (Hyclone, cat # SH30071.03) and 100 U/mL penicillin/streptomycin (Invitrogen, cat # 15140). Cell line identities were confirmed by short tandem repeat profiling.^77^ Cell cultures were routinely tested for *Mycoplasma* contamination using the MycoAlert PLUS *Mycoplasma* Detection Kit (Lonza Bioscience, cat # LT07) according to manufacturer protocol. Cell counts were measured by Countess automated cell counter (Invitrogen) and 0.4% trypan blue solution (Invitrogen, cat # T10282). The following reagent was deposited by the Centers for Disease Control and Prevention and obtained through BEI Resources, NIAID, NIH: SARS-Related Coronavirus 2, isolates USA-WA1/2020 (NR-53872).

### Ivermectin quality control

The ^1^H NMR (400 MHz) and ^13^C NMR (101 MHz) spectra were recorded on a Bruker spectrometer. Chemical shifts are reported in ppm using the solvent peak as an internal standard. Analytical analysis was performed on an Agilent LC-MS (Agilent Technologies, Santa Clara, CA) as follows: a 7 min gradient of 4% to 100% acetonitrile (containing 0.025% trifluoroacetic acid, v/v) in water (containing 0.05% trifluoroacetic acid, v/v) was used with an 8 min run time at a flow rate of 1 mL/min. A Phenomenex Luna C18 column (3 μm, 3 x 100 mm) was used at a temperature of 50 °C. Mass determination was performed using an Agilent 6130 mass spectrometer with electrospray ionization in the positive mode. Purity determination was performed using an Agilent Diode Array Detector and an Agilent Evaporative Light Scattering Detector. NMR and UPLC-MS data were processed and analyzed using Mnova version 12 (Mestrelab).

### *C. elegans* viability assay

The activity of the primary ivermectin sample (**1a**) was assessed using a previously described *C. elegans* quantitative high-throughput screening assay for nematode viability.^20^ Briefly, test compounds were arrayed in 11-pt, 1:3 or 22-pt, 1:2 intra-plate titrations. Compounds were tested at a high concentration of 0.1-25 μM for final assay titrations spanning 0.20 pM – 333 nM to 39.7 pM – 83.3 μM. Levamisole HCl control was arrayed in an 8-pt 1:3 titration spanning 15.2 nM to 33.3 μM final assay concentration range. All compounds were plated in 384-well black, μClear microtiter plates (Greiner, cat # 781096) with an Echo 555 acoustic dispenser. *E. coli* ghosts media were diluted to OD_600_ 0.81 in S medium (IPM, cat # 11006-505) and manually added at 30 μL per well to assay plates with an Integra multichannel pipette. *C. elegans* strain PE254 (acquired from the University of Minnesota *Caenorhabditis* Genetics Center) ubiquitously expressing GFP were grown on nematode growth media plates (Teknova, cat # N1098) with OP50 *E. coli* as a nutrient source. Worms were collected from plates, washed, bleached, and axenized eggs were hatched overnight in M9 buffer at 20 ^°^C. Synchronized L1 worms were diluted in S medium and plated at ten worms per 30 μL per well with an Integra multichannel pipette. Plates were sealed with Breath-Easy sealing membranes (cat # Z380059) and incubated at 20 ^°^C for 7 d. Worm viability was assessed as GFP area per well measured on an Acumen laser scanning cytometer. Data was normalized to 41.7 μM levamisole HCl intra-plate control as –100% activity and DMSO neutral control as 0% activity.

### NCATS OpenData COVID-19 portal analyses

The following biochemical assay results were accessed from the public domain (https://opendata.ncats.nih.gov/covid19): Spike-ACE2 protein-protein interaction (AID 1), Spike-ACE2 protein-protein interaction TruHits counter-screen (AID 2), ACE2 enzymatic activity (AID 6), TMPRSS2 enzymatic activity (AID 8), SARS-CoV-2 3CL protease activity (AID 9), and SARS-CoV-2 RNA-dependent RNA-polymerase activity (AID 10). Detailed experimental protocols for each assay have been disclosed (https://opendata.ncats.nih.gov/covid19/assays).

### SARS-CoV-2 FLuc assay

The antiviral activity of compounds were assessed by a live SARS-CoV-2 replication assay in A549-ACE2 host cells using an engineered SARS-CoV-2 WA-1 lineage virus containing an integrated firefly luciferase reporter (FLuc, provided by Pei-Yong Shi, UTMB).^78^ Experiments were performed in a BSL-3 facility. Briefly, 20 nL/well of compounds or DMSO controls were spotted onto black, clear bottom, sterile, tissue culture-treated 1536-well assay microplates (Aurora, cat # EBA1410000A) by Echo 655 acoustic dispenser. A MultiDrop (Thermo Fisher) and small-volume metal tip dispensing cassette (Thermo Fisher, cat # 24073295) was then used to dispense 4 μL of A549-ACE2 cells (1600 cells/well) in DMEM with 2% FBS (without antibiotics) to all wells, followed by the addition of 1 μL cell-free media into control microplate wells, followed by the addition of 1 μL of SARS-CoV-2 (USA_WA1/2020, multiplicity of infection = 0.2) suspended in media to the remaining microplate wells. Microplates were then covered with vented cell culture-compatible metal lids (Wako), and the assay plates were then incubated for 48 h at 37 °C, 5% CO_2_, and 90% humidity. After incubation, 2 μL/well of One-Glo detection reagent (Promega, cat # E6120) was added to all microplate wells *via* Multidrop, followed by 5 min incubation at RT. Luminescence signal was measured on a BMG PheraStar (1536-well aperture, 0.2 sec exposure). Raw data were normalized to the neutral control (cells infected with virus + DMSO, set as 0%) and positive control (cells without virus added, set as – 100%) for each plate. The cell viability counter-screen was performed similarly, except that the addition of virus was omitted. After 48 h compound treatment, 2 μL of CellTiter-Glo reagent (Promega, cat # G9683) was dispensed to each microplate well *via* BioRAPTR FRD. Cell viability was then measured using a ViewLux μHTS Microplate Imager (PerkinElmer). Raw data were normalized to the neutral control (cells + DMSO, set as 0%) and positive control (media + DMSO, set as –100%) for each plate. Data are mean ± SD of five experimental replicates.

### High-content SARS-CoV-2 assay

Antiviral activity of compounds were also assessed using a live SARS-CoV-2 virus high-content imaging assay utilizing wild-type viral isolates in A549-ACE2-TMPRSS2 host cells. Experiments were performed in a BSL-3 facility. Briefly, 40 nL/well of compounds or DMSO controls were spotted onto black, optically clear bottom, sterile, tissue culture-treated 384-well assay microplates (PerkinElmer Phenoplate, cat # 6057308) by Echo 655 acoustic dispenser. A MultiDrop was then used to dispense 15 μL of A549-ACE2-TMPRSS2 cells (4,500 cells/well) in DMEM with 2% FBS (without antibiotics) to all wells, followed by the addition of 5 μL cell-free media into control microplate wells, followed by the addition of 5 μL of SARS-CoV-2 strain virus (multiplicity of infection = 0.08) suspended in media to the remaining microplate wells. Microplates were then incubated for 48 h at 37 °C, 5% CO_2_, and 90% humidity. After incubation, media was evacuated using a BlueWasher (BlueCatBio), and 20 μL/well of 4% paraformaldehyde was dispensed using the BlueWasher. After 20 min of room-temperature incubation, wells were washed twice with 20 μL/well of PBS. Wells were subsequently evacuated and incubated overnight at 4 °C with 20 μL/well of primary antibody (anti-SARS-CoV-2 nucleocapsid antibody, SinoBiological, cat # 40143-R040 mAB, 1:2000 dilution) diluted in PBTG (PBS with 1% BSA, 5% normal goat serum and 0.1% Triton X-100). After this incubation, wells were evacuated using the BlueWasher. Next, 20 μL of a solution containing secondary antibody (Cell Signaling Technologies, cat # 4414S, anti-Rb mAb-Alexa 647 conjugate, 1:2000 dilution) and 1:10000 Hoechst 33342 diluted in PBTG was dispensed to each well, followed by incubation at room temperature for 30 min. Plates were then washed four times with 20 μL/well PBS using the BlueWasher.

High-content imaging of the wells were then acquired using a PerkinElmer Opera Phenix. Confocal digital micrograph images were captured for two fluorescence channels per sample: excitation/emission 405/456 nm (nuclei) and 640/704 nm (nucleocapsid), using a 10X air objective with digital camera binning set at 2X, resulting in a final in image x-y pixel size of 1.196 microns. For each sample well in the microplate 9X adjacent x-y image fields were captured. Digital 16-bit gray scale images were transferred to the PerkinElmer Columbus v2.9.1 digital image analysis platform. A fully automated Columbus image analysis routine was applied to segment the nuclear and cytoplasmic regions of each individual cell. Intensity measurements in the 640 nm fluorescence channel in each whole cell region were extracted and digitally stored. To apply a consistent 640 nm intensity threshold for identification of ‘viral positive cells’ across all experiments, a customized Matlab script was created and applied to the single cell-level measurements. This fully automated script calculates the mean 640 nm cellular intensity and cellular standard deviation of this signal in all negative control wells (those wells that were not infected but were stained for virus, providing measurement of background non-specific fluorescence) on each plate. The threshold is defined as the mean negative control intensity plus 4x the negative control standard deviation. The script applies this threshold to all sample wells and calculates ‘percent viral positive cells’ per well.

### Cellular health assays

A general procedure for cellular health assays is as follows: For cellular health assays, 4 μL of cells (cell seeding density 500 cells/μL; 2000 cells total per well) were plated into white 1536-well microplates (Greiner, cat # 789173-F) *via* Multidrop dispenser (Thermo Fisher) and small-volume metal tip dispensing cassette (Thermo Fisher, cat # 24073295). For some assay formats, 4 μL of cell-free complete media were dispensed to control columns to serve as background controls. Cells were then covered with vented cell culture-compatible metal lids (Wako), and then incubated for 24 h at 37 °C with 5% CO_2_ and 95% humidity. Test compounds or DMSO controls were then acoustically dispensed to respective wells of each microplate at 20 nL/well *via* Echo 655 in 12-point, 1:2 titrations. Final DMSO concentration was 0.50%. Cells were incubated for 6, 12, 24, 48, or 72 h at 37 °C with 5% CO_2_ and 95% humidity. Each time point consisted of a single microplate containing all technical replicates and controls. After this compound treatment interval, reagents were dispensed to the microplate wells *via* BioRAPTR FRD. Microplates were then incubated per manufacturer protocols. Microplates were then measured with either a ViewLux 1430 Ultra HTS (PerkinElmer; luminescence) or Spark (Tecan; fluorescence intensity). Staurosporine and digitonin were used as positive cytotoxic control compounds. Remdesivir was used as a negative cytotoxic control compound. Data are mean ± SD of four intra-plate technical replicates.

A summary of the important assay-specific considerations includes:

- ***Cell viability assay (CellTiter-Glo, ATP content).*** CellTiter-Glo Luminescent Cell Viability Assay (“CTG”; Promega, cat # G7572) was used to quantify cellular ATP according to manufacturer protocol. Microplates were incubated under reduced lighting at ambient temperature for 10 min after reagent addition. Luminescence was quantified with the following optical settings: exposure = 1 sec, gain = high, speed = slow, binning = 2X. Data were normalized to negative control (DMSO, 100% viability) and positive control (125 μM digitonin, 0% viability).
- ***Cell viability assay (resazurin, reducing ability).*** CellTiter-Blue Cell Viability Assay (Promega, cat # G8080) was used to quantify cellular reducing ability according to manufacturer protocol. Microplates were incubated at 37 °C with 5% CO_2_ and 95% humidity for 1 h after adding 1 μL rezasurin reagent per microplate well *via* BioRAPTR FRD. Next, 2 μL stop solution was added to each microplate well *via* BioRAPTR FRD. Microplates were incubated under reduced lighting at ambient temperature for 10 min. Fluorescence intensity was quantified with the following optical settings: excitation 560 nm, emission 590 nm, 10 nm bandwidth filters, monochromator, gain = extended dynamic range mode. Data were normalized to negative control (DMSO, 100% viability) and positive control (125 μM digitonin, 0% viability).
- ***Cell viability assay (RealTime-Glo, reducing ability).*** RealTime-Glo MT Cell Viability Assay (Promega, cat # G9711) was used to quantify cellular reducing ability according to manufacturer protocol. Microplates were incubated under reduced lighting at ambient temperature for 10 min after the addition of 4 μL 2X reagent solution to each microplate well *via* BioRAPTR FRD. Luminescence was quantified with the following optical settings: exposure = 10 sec, gain = high, speed = slow, binning = 2X. Data were normalized to negative control (DMSO, 100% viability) and positive control (125 μM stippled lines at 0% denote no effect relative to control; stippled lines at –100% denotes complete cell killing or complete virus reduction in, 0% viability).
- ***Cytotoxicity assay (LDH release).*** CytoTox-ONE Homogeneous Membrane Integrity Assay (Promega, cat # G7890) was used to quantify cytotoxicity by LDH release according to manufacturer protocol. At the indicated time points, microplates were removed from cell culture incubators and allowed to cool at ambient temperature for 30 min. Next, 500 nL of lysis reagent was manually dispensed to control columns *via* multichannel pipette (to serve as control for maximum possible LDH release). Next, 4 μL 2X reagent solution was dispensed to each microplate well *via* BioRAPTR FRD, followed by microplate incubation under reduced lighting at ambient temperature for 10 min. Fluorescence intensity was quantified with the following optical settings: excitation 560 nm, emission 590 nm, 10 nm bandwidth filters, monochromator, gain = extended dynamic range mode. Data were normalized to negative control (DMSO, 0% relative cytotoxicity) and positive control (DMSO plus lysis solution, 100% relative cytotoxicity).
- ***Cytotoxicity assay (CellTox Green, membrane permeability).*** CellTox Green Cytotoxicity Assay (Promega, cat # G8741) was used to quantify cytotoxicity according to manufacturer protocol. At the indicated time points, 500 nL of lysis reagent was manually dispensed to control columns *via* multichannel pipette (to serve as control for maximum nuclear dye binding). Microplates were then incubated under reduced lighting at ambient temperature for 30 min. Next, 4 μL 2X reagent solution was dispensed to each microplate well *via* BioRAPTR FRD, followed by microplate incubation under reduced lighting at ambient temperature for 10 min. Fluorescence intensity was quantified with the following optical settings: excitation 490 nm, emission 530 nm, 10 nm bandwidth filters, monochromator, gain = optimal. Data were processed by background signal subtraction (cell-free media plus reagent). Data were calculated as percentage relative to negative control (DMSO).
- ***Apoptosis assay (caspase-3/7).*** Caspase-Glo 3/7 Assay System (Promega, cat # G8091) was used to quantify activated cellular caspase 3/7 per manufacturer protocol. Microplates were incubated under reduced lighting at ambient temperature for 30 min after adding 4 μL 2X reagent solution to each microplate well *via* BioRAPTR FRD. Luminescence was quantified with the following optical settings: exposure = 1 sec, gain = high, speed = slow, binning = 2X. Data were processed by background signal subtraction (cell-free media plus reagent). Data were calculated as percentage relative to negative control (DMSO).
- ***Apoptosis assay (annexin V).*** RealTime-Glo Annexin V Apoptosis Assay (Promega, cat # JA1000) was used to quantify annexin V binding according to manufacturer protocol. Microplates were incubated under reduced lighting at ambient temperature for 10 min after adding 4 μL 2X reagent solution (prewarmed, in complete media) to each microplate well *via* BioRAPTR FRD. Luminescence was quantified with the following optical settings: exposure = 10 sec, gain = high, speed = slow, binning = 2X. Data were processed by background signal subtraction (cell-free media plus reagent). Data were calculated as percentage relative to negative control (DMSO).
- ***Cellular media assay (FBS titration).*** The effect of FBS on compound-mediated cellular injury was assessed by the aforementioned CTG protocol with minor modifications. For FBS titration experiments, cells were harvested and then resuspended in FBS-free media (DMEM, 0% FBS, 100 U/mL penicillin/streptomycin). Seeding solutions were prepared by the addition of the appropriate volumes of 100% FBS or DPBS to generate seeding solutions containing 0, 2, 5, 10, or 20% FBS (v/v). Cells were then dispensed into microplate columns *via* Multidrop dispenser. Each microplate contained all experimental conditions and technical replicates. Data were normalized to negative control (DMSO, 100% viability) and positive control (125 μM digitonin, 0% viability) for each FBS concentration.
- ***Cellular media assay (BSA titration).*** The effect of BSA on compound-mediated cellular injury was assessed by the aforementioned CTG protocol with minor modifications. For BSA titration experiments, cells were harvested and then resuspended in compete media (DMEM, 2% FBS, 100 U/mL penicillin/streptomycin). Seeding solutions were prepared by the addition of the appropriate volumes of sterile 35% BSA in DPBS (Sigma, cat # A7979) or DPBS to generate seeding solutions containing 0, 37, 111, 333, or 1000 μM supplemental BSA. Cells were then dispensed into microplate columns *via* Multidrop dispenser. Each microplate contained all experimental conditions and technical replicates. Data were normalized to negative control (DMSO, 100% viability) and positive control (125 μM digitonin, 0% viability) for each BSA concentration.

### Broad PRISM and NCI60 analyses

Broad PRISM data were accessed from the Broad DepMap (https://depmap.org/portal/) for the following compounds: IVM, BRD-K85554912-001-06-3, dataset 19Q4; selamectin, BRD-A58564983-001-04-6, dataset 19Q4; and eprinomectin, BRD-A91366704-001-01-9, dataset 19Q4. The Broad PRISM methodology has been previously described.^79^ NCI60 tumor profiling data were accessed from the public domain (https://dtp.cancer.gov) for the following compounds: abamectin, NSC-758202; doramectin NSC-760342; emamectin, NSC-781687; and selamectin, NSC-758615. The NCI60 methodology has been previously described (https://dtp.cancer.gov/discovery_development/nci-60/methodology.htm).

### Alpha technology counter-screens

Compound-mediated interference with the Alpha homogenous proximity technology was assessed as described in the NCATS *Assay Guidance Manual*.^27^ The buffer composition, incubation times, reagent concentrations, and order of reagent addition were designed to model a previous report.^7^ Most compounds were tested at 14-point, 1:2 titrations with a maximum final concentration of 500 µM (except for biotin: 100 µM maximum final concentration, 1:3 titration). A summary of the important assay-specific considerations include:

- ***UV/Vis absorption spectra.*** To a black, optically clear, flat bottom, cyclic olefin 384-well microplate (CellCarrier Ultra, PerkinElmer, cat # 6057300), 125 µL buffer containing 1X PBS (pH 7.4; Gibco, cat # 10010023), 0.1% BSA (Sigma, cat # A7979), and ± 0.1% Tween-20 (v/v; Sigma, cat # P1379) was added to each well *via* multichannel pipette. Next, 5 µL of test compounds or DMSO controls were dispensed to respective microplate wells *via* multichannel pipette. Final DMSO concentration was 2.0%. Absorption spectra were collected from 250 to 800 nm at 1 nm intervals using a Tecan Spark. Data are mean ± SD of three intra-plate technical replicates.
- ***AlphaScreen with biotin-His6 substrate.*** AlphaScreen Histidine (Nickel Chelate) Detection Kit (PerkinElmer, cat # 6760619C) and biotinylated-His6 substrate (PerkinElmer, cat # 6760303M) were used according to manufacturer instructions with minor modifications. First, 100 nL of test compounds or DMSO controls were acoustically dispensed to grey, flat bottom, non-sterile, non-tissue culture-treated 1536-well microplates (Aurora, cat # EWB010000A) at *via* Echo 655. Next, 2.5 μL of nickel chelate acceptor beads (2X solution, 40 µg/mL) in buffer (1X PBS, pH 7.4, 0.1% BSA, 10 nM biotin His6, ± 0.1% Tween-20 v/v) was added to microplate wells *via* BioRAPTR FRD. Microplates were covered with non-vented metal lids (Wako) and incubated at ambient temperature under reduced lighting for 90 min. Next, 2.5 µL of streptavidin donor beads (2X solution, 40 µg/mL) in buffer (1X PBS, pH 7.4, 0.1% BSA, ± 0.1% Tween-20 v/v) was added to microplate wells *via* BioRAPTR FRD. Microplates were incubated at ambient temperature under reduced lighting for 2 h. Final DMSO concentration was 2.0%. Fluorescence intensity was quantified using a PHERAstar FSX (BMG Labtech) with the following optical settings: excitation 680 nm, emission 570 nm (AlphaScreen optical module), gain = 2000, speed = maximum precision, cross-talk correction read order. Data were normalized to negative control (DMSO, 100% expected signal) and positive control (DMSO, acceptor beads only, 0% expected signal). Data are mean ± SD of four intra-plate technical replicates. In a modification of this protocol (prebound beads), compounds or DMSO were acoustically dispensed to microplate wells 30 min after the addition of donor beads to acceptor beads, followed by incubation at ambient temperature under reduced lighting for 30 min before measuring fluorescence.
- ***TruHits AlphaScreen counter-screen.*** AlphaScreen TruHits Kit (PerkinElmer, cat # 6760627D) was used according to manufacturer instructions with minor modifications. First, 100 nL of test compounds or DMSO controls were acoustically dispensed to grey, flat bottom, non-sterile, non-tissue culture-treated 1536-well microplates (Aurora, cat # EWB010000A) at *via* Echo 655. Next, 2.5 µL of biotinylated acceptor beads (2X solution, 10 µg/mL) in buffer (1X PBS, pH 7.4, 0.1% BSA, ± 0.1% Tween-20 v/v) was added to microplate wells *via* BioRAPTR FRD. Microplates were covered with non-vented metal lids (Wako) and incubated at ambient temperature under reduced lighting for 90 min. Next, 2.5 µL of streptavidin donor beads (2X solution, 20 µg/mL) in buffer (1X PBS, pH 7.4, 0.1% BSA, ± 0.1% Tween-20 v/v) was added to microplate wells *via* BioRAPTR FRD. Microplates were incubated at ambient temperature under reduced lighting for 2 h. Final DMSO concentration was 2.0%. Fluorescence intensity was quantified using a PHERAstar FSX (BMG Labtech) with the following optical settings: excitation 680 nm, emission 570 nm (AlphaScreen optical module), gain = 2000, speed = maximum precision, cross-talk correction read order. Data were normalized to negative control (DMSO, 100% expected signal) and positive control (DMSO, acceptor beads only, 0% expected signal). Data are mean ± SD of four intra-plate technical replicates. In a modification of this protocol (prebound beads), compounds or DMSO were acoustically dispensed to microplate wells 30 min after the addition of donor beads to acceptor beads, followed by incubation at ambient temperature under reduced lighting for 30 min before measuring fluorescence.
- ***TruHits AlphaLISA counter-screen.*** AlphaLISA TruHits Kit (PerkinElmer, cat # AL900) was used according to manufacturer instructions. The counter-screen was performed similarly to the AlphaScreen TruHits counter-screen with the noted modifications. Fluorescence intensity was quantified with the following optical settings: excitation 680 nm, emission 615 nm (AlphaLISA optical module).
- ***OmniBeads counter-screen.*** AlphaScreen OmniBeads (PerkinElmer, cat # 6760626D) were used according to manufacturer instructions with minor modifications. First, 100 nL of test compounds or DMSO controls were acoustically dispensed to grey, flat bottom, non-sterile, non-tissue culture-treated 1536-well microplates (Aurora, cat # EWB010000A) at *via* Echo 655. Next, 5 µL of Omnibeads beads (1X solution, 20 µg/mL) in buffer (1X PBS, pH 7.4, 0.1% BSA, ± 0.1% Tween-20 v/v) was added to microplate wells *via* BioRAPTR FRD. Microplates were covered with non-vented metal lids (Wako) and incubated at ambient temperature under reduced lighting for 90 min. Final DMSO concentration was 2.0%. Fluorescence intensity was quantified using a PHERAstar FSX (BMG Labtech) with the following optical settings: excitation 680 nm, emission 570 nm (AlphaScreen filter module), gain = 2000, speed = maximum precision, cross-talk correction read order. Data were normalized to negative control (DMSO, 100% expected signal) and positive control (DMSO, buffer only, 0% expected signal). Data are mean ± SD of four intra-plate technical replicates.

### Aggregation and phospholipidosis counter-screens

IVM was assessed for aggregation using an AmpC β-lactamase counter-screen, a malate dehydrogenase counter-screen, and dynamic light scattering as previously described.^80, 81^ Drug-induced phospholipidosis was assessed using an NBD-PE high-content assay as previously described.^31^

### GPCR profiling

The binding of IVM (**1a**) to GPCRs was profiled by radiolabeled binding assays by the NIMH Psychoactive Drug Screening Program.^38^ In primary binding testing, IVM was tested at 10 μM final concentrations with four intra-plate technical replicates. In secondary binding to determine *K*_i_ values, IVM (was tested at 12 concentrations with half-log serial dilutions (10 μM to 0.01 nM final concentrations) with three inter-plate technical replicates in three to four independent experiments. The functional effect of IVM on GPCR agonist and antagonist activity in HEK293 cells was performed by the PDSP using two formats: a Tango β-arrestin assay^40, 82^ and a GloSensor cAMP assay.^41^ The functional effect of IVM on neurotransmitter transporter proteins (dopamine, norepinephrine, serotonin) in HEK293 cells was profiled by the PDSP using a fluorescent dye reuptake assay. In these secondary functional assays, IVM was tested at 16 concentrations with half-log serial dilutions (usually 30 μM to 1 pM final concentrations) with three to four inter-plate technical replicates in three to four independent experiments. Detailed descriptions of each assay are also available in the PDSP Assay Protocols (https://pdsp.unc.edu/pdspweb/?site=assays). Reference control compounds were included in each PDSP experiment.

The functional effect of IVM on GPCR activity was also performed using an orthogonal β-arrestin reporter assay (gpcrMAX, Eurofins).^39^ A total of 168 unique GPCRs were tested in both agonist and antagonist modes at 10 μM IVM (**1a**) final concentrations. Samples were tested in duplicate along with vendor-provided reference compounds for quality control.

### PubChem analyses

Data was downloaded from PubChem (https://pubchem.ncbi.nlm.nih.gov, data accessed 25 July 2023). The relevant PubChem CIDs for ivermectin (3085416, 6321424, 11957587, 45114068, and 73265241) were identified by keyword searches (“ivermectin”, “NCGC00094047”) and manually verified for the correct chemical structure and association with primary screening data. Bioactivity records annotated by PubChem as “literature-derived” or “summary” were excluded. Duplicate bioassay records were excluded unless independent samples were tested. Secondary assays were excluded unless they were performed as part of a whole-deck screen. Validation HTS experiments using the LOPAC were included. Each bioassay was annotated as “biochemical” (cell-free) or “cellular” (cell-based, includes bacteria, yeast, and *in vivo* assays) by manually inspecting the deposited PubChem protocols, with the analyst blinded to the activity status during the annotation process. The activity summary provided by PubChem was used to determine activity (active, inactive, inconclusive, unspecified, probe). Data were analyzed by Chi-square test to assess if the activity classifications were equally distributed between biochemical and cellular assays.

### Membrane bilayer assays

Drug-induced membrane bilayer perturbations were quantified using both fluorescence quench and single-channel methods.

- ***Fluorescence quench experiments***. These experiments were done as previously described.^49^ Briefly, 1,2-dierucoyl-sn-glycero-3-phosphocholine (DC_22:1_PC; Avanti Polar Lipids, cat # 850398): the natural mixture of linear gramicidins, gramicidin D (Sigma-Aldrich, cat # G5002) were mixed in a 2000:1 molar ratio in chloroform was dried and rehydrated in fluorophore (ThermoFisher, cat # A350) buffer (25 mM ANTS, 100 mM NaNO3, 10 mM HEPES, pH 7). Large unilamellar vesicles were prepared by passing the rehydrated lipid suspension through 0.1-µm membrane using an Avanti Mini-extruder, and extravesicular ANTS was removed using a PD-10 desalting column (GE Healthcare, cat # 17085101). The rate of ANTS fluorescence quenching by the gA channel-permeant Tl^+^ was recorded with an Applied Photophysics SX.20 (Leatherhead, UK) stopped-flow spectrofluorometer with dead time < 2 ms. Samples were excited using 360 nm LED and the fluorescence signal above 455 nm was recorded. Vesicles were incubated with the compound for 10 min at 25 °C before measuring the quenching rate. Nine mixing reactions were done with each sample. Each reaction was analyzed separately using MATLAB v7.9 (The MathWorks Inc.). Quench rates were determined by fitting the first 1 s of the fluorescence time course for the individual reactions with the modified stretched exponential^21^:

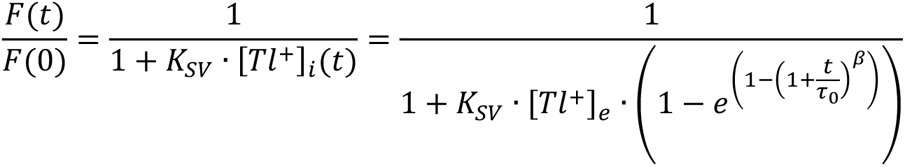

The rate was evaluated using 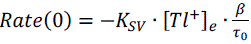, where K_SV_ is the Stern-Volmer constant, [Tl^+^]_e_ is the external Tl^+^ concentration, where τ_0_ (τ_0_ > 0) is a parameter with units of time, and β (0 < β ≤1) a parameter reflecting the vesicle dispersity as reflected in the distribution of quench rates in different vesicle populations. Quench rates measured in the presence of the tested compound (*Rate*) were normalized to rates in the absence of the compound (*Rate*_control_) to yield *NormRate = Rate*/*Rate*_control_). The experiments were done in duplicate and reported as mean ± SD.

- ***Single-channel experiments***. These experiments were done using the bilayer punch method has been described previously.^52^ In brief, a bilayer composed of 1,2-dioleoyl-*sn*-glycero-3-phosphocholine (DC_18:1_PC, Avanti Polar Lipids, cat # 850375) suspended in *n*-decane was formed across a 1–1.5 mm hole in a Teflon partition and doped with gA analogs of different lengths and helix sense: the 15-amino acid residue [Ala^1^]gramicidin A (AgA(15)) with the sequence f-AGA<u>L</u>A<u>V</u>V<u>V</u>W<u>L</u>W<u>L</u>W<u>L</u>W-e, where f denotes formyl and e ethanolamine and the underlined residues are D-amino acids (glycine is achiral), which forms right-handed channels; and the chain-shortened 13-amino acid residue enantiomer des-Val^1^,Gly^2^-gramicidin A– (gA–(13)) with the sequence f-<u>A</u>L<u>A</u>V<u>V</u>V<u>W</u>L<u>W</u>L<u>W</u>L<u>W</u>-e, where the underlined residues are D-amino acids, which forms left-handed channels. (The total AgA(15) concentration in the system was a few pM in the DC_18:1_PC experiments; the concentration of gA^−^(13) was about 10-fold higher.) Punch electrodes, bent 90°, with a 20-40 µm opening were used to isolate small membrane patches and record electrical activity. The measurements were done in 1 M NaCl (plus 10 mM HEPES, pH 7) at 25 ± 1 °C, at an applied potential of ± 200 mV. The current signal was recorded and amplified using a Dagan 3900A (Minneapolis, MN) patch clamp, filtered at 5 kHz, digitized, and sampled at 20 kHz by a PC/AT compatible computer, and filtered at 200 – 500 Hz. Single-channel events were detected using a transition-based algorithm, and single-channel lifetimes were determined as described previously.^83^ The average lifetimes were determined by fitting the results with single exponential distributions:

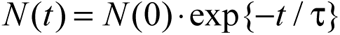

where τ denotes the average channel lifetime and *N*(*t*) the number of channels with durations longer than *t*, which was fit to the lifetime distributions using the non-linear least-squares fitting routine in Origin 8.1 (OriginLab). The single-channel conductance and lifetime results are reported is mean ± SD using the weighted mean and SD of results for three to six experiments, where the weighted mean (mean_W_) and SD (SD_W_) are calculated as:

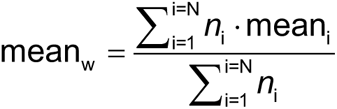

and

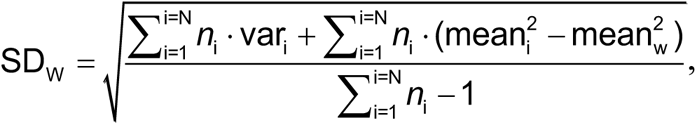

where *n*_i_, mean_i_ and var_i_ denote number of events, mean and variance of experiment *i*.

### Sodium channel assay

Endogenous TTX-sensitive Na^+^ currents were recorded from the mammalian neuronal cell line ND7/23 as previously described.^57^ In brief, cells were grown on 12-mm coverslips and whole cell voltage-clamp recordings were performed at room temperature (22-24 °C) with a patch-clamp amplifier (Axopatch 200B, Molecular Devices) using a 5 kHz low-pass filter and a sampling rate of 20 kHz. The external bath solution contained (in mM): 130 NaCl, 10 HEPES, 3.25 KCl, 2 MgCl_2_, 2 CaCl_2_, 20 TEACl, and 5 D-glucose, adjusted to pH 7.4 (with NaOH), and 310 mOsm/kg H_2_O (with sucrose). Recording pipettes were pulled from borosilicate glass capillaries (Sutter Instruments, Novato, CA) using a P-1000 puller (Sutter Instruments). Pipettes had a tip resistance of 1.5-2.5 MΩ when filled with following pipette solution (in mM): 120 CsF, 10 NaCl, 10 HEPES, 10 EGTA, 10 TEACl, 1 CaCl_2_, and 1 MgCl_2_, adjusted to pH 7.3 (with CsOH), and 310 mOsm/kg H_2_O (with sucrose). Access resistance was further decreased using 70-80% series resistance compensation. Liquid-junction potentials were not corrected; capacitative current transients were electronically cancelled with the internal amplifier circuitry. A stock solution of IVM was prepared in DMSO and further diluted to the final working concentration with external bath solution prior to the experiment. Control solutions had the same amount of DMSO as the drug solutions (< 0.1%). Cells were superfused using a pressurized perfusion system (ALA Scientific Instruments) with a 200 µm-diameter perfusion pipette positioned in close proximity of the cell. The holding potential (*V*_h_) was –80 mV unless otherwise stated. Voltage-dependent inhibition of peak inward Na^+^ current (*I*_Na_) was determined using a 10 ms test pulse to 0 mV preceded by a 300 ms prepulse alternating between either –130 mV (V_0_) or the voltage of half-maximal inactivation (V_½_ = –69 ± 3 mV) applied every 5 s. *V*_½_ was determined for each individual cell using the double-pulse protocol for steady-state inactivation described below. Steady-state fast inactivation (*h*_∞_) was measured using a double-pulse protocol with 300 ms prepulses ranging from –130 to –30 mV in 10 mV steps, followed by 10 ms test pulses to 0 mV. Peak currents during the test pulse were normalized to the maximal current (*I*/*I*_max_, where *I*_max_ is the maximal current that is elicited at the test potential), and plotted vs. the prepulse potential (*V*_m_), and fitted with a Boltzmann function^52^:

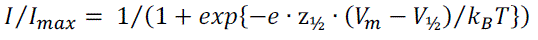

where z_½_ and *V*_½_ denote the apparent gating valence and potential for half-maximal inactivation, respectively.

### Statistics and reproducibility

No statistical method was used to predetermine sample size. Select replicates were excluded from analyses due to technical factors such as suspected automated liquid dispensing errors (noted in Source Data file). The experiments were not randomized. Except for the experiments summarized in **Supplementary Table 1**, the investigators were not blinded to allocation during experiments and outcome assessment. The sample size of at least three technical replicates were chosen based on internal experience with qHTS. For most experiments, the number of technical replicates (usually three) are sufficient to determine significant differences between compounds within a high-throughput experiment.

### Data analyses and figure preparation

All graphical data are expressed as individual replicates or mean ± standard deviation (SD) unless stated otherwise. Graphing and statistical analyses were performed using GraphPad Prism (version 9.0.0) or OriginLab Origin (versions 8.1 and 8.5 for the gramicidin experiments). Concentration-response curves were generated using the sigmoidal dose-response variable slope four-parameter equation in GraphPad Prism. Outliers and concentration-response curves with poor fits were excluded from certain analyses, and in such cases are noted in the Source Data files. Final figures were prepared in Adobe Illustrator (version 25.0.1). Statistical differences were determined by: ordinary one-way ANOVA adjusted for multiple comparisons (**Figure 2C**, **Supplementary** Figure 2A) and two-tailed Mann-Whitney test (**Table 2**).

## Data availability

All relevant data are available from the authors without restriction. Source data are provided as Source Data files.

## Preprint status

This manuscript and supplementary materials have been uploaded to bioRxiv.

## Supplementary materials

Supplementary Information: file containing Supplementary Figures 1-8, Supplementary Tables 1-4, Supplementary Notes, and Supplementary References (PDF). Source Data 1: processed data for Figures 1-6, Tables 1-2, and Supplementary Figures 1-8 (XLS). Source Data 2: unprocessed NMR and UPLC-MS files from Supplementary Figure 8 (ZIP).

## ORCID iDs

RE: 0000-0003-3580-380X

RR: 0000-0003-3709-5851

KFH: 0000-0002-8614-0578

XPH: 0000-0002-2585-653X

PD: 0000-0002-1349-4885

TCV: 0009-0002-8750-2478

SR: 0000-0001-5802-5970

JHS: 0000-0001-9872-942X

ADW: 0000-0003-2180-2136

HCH: 0000-0002-6043-5482

BLR: 0000-0002-0561-6520

JI: 0000-0002-7332-5717

OSA: 0000-0002-3026-6710

JLD: 0000-0003-4151-9944

## Author contributions (CRediT)

Conceptualization: OSA, JI, JLD. Data curation: OSA, TCV, JLD. Formal analysis: RR, KFH, OSA, JLD. Investigation: RR, RTE, KFH, XPH, PD, TYC, JHS, ADW, SR, JLD. Methodology: OSA, JLD. Supervision: XPH, BLR, HCH, JI, OSA, JLD. Validation: JLD. Visualization: OSA, JLD. Writing, original draft: JLD. Writing, review and editing: all authors.

## Acknowledgements

The authors acknowledge: Isabella Glenn and Dr. Brian Shoichet for assistance with aggregation and drug-induced phospholipidosis counter-screens (sponsored by NIH R35GM122481); Paul Shinn, Zina Itkin, and NCATS Compound Management for assistance with sample management and quality control; Erin Oliphant for tissue culture support; Adam Yasgar for technical assistance; Dr. Quinlin M. Hanson for helpful discussions; the NCATS COVID-19 OpenData team; and E. Ashley Hobart for assistance with the gramicidin single-channel experiments. This research was supported in part by the Intramural Research Program of the National Center for Advancing Translational Sciences, National Institutes of Health under project 1ZIATR000052 (JI) and the A Specialized Platform for Innovative Research Exploration (ASPIRE) program. OSA and RR were supported by NIH grant GM021342-45. KH and HCH were supported by NIH grant GM058055-24. GPCR profiling was provided by the National Institute of Mental Health’s Psychoactive Drug Screening Program, contract # HHSN-271-2018-00023-C (NIMH PDSP). The content of this publication does not necessarily reflect the views or policies of the Department of Health and Human Services, nor does mention of trade names, commercial products, or organizations imply endorsement by the U.S. Government.

## Conflicting interests

JLD is currently an employee of Agios Pharmaceuticals. RTE is currently an employee of Novartis Pharmaceuticals. The remaining authors hereby declare no conflicting interests pertaining to the material in this manuscript.

