## Supplementary Information for "Nonspecific membrane bilayer perturbations by ivermectin underlie SARS-CoV-2 *in vitro* activity"

### Table of contents

Supplementary figures

S1. Cellular health control compounds S2
S2. Cellular health ivermectin analogs S3
S3. Complete media content can modulate effect of ivermectin on cell viability S4
S4. Alpha technology interference profiles of reference compounds S5
S5. Ivermectin is unlikely to interfere with biological assays by colloidal aggregation or drug-induced phospholipidosis S7
S6. Ivermectin and chemical analogs do not inhibit several SARS-CoV-2-related targets in cell-free assays S7
S7. Ivermectin does not inhibit recombinant luciferases S8
S8. Quality control of primary ivermectin sample S9

Supplementary tables

S1. Summary of ivermectin target activities S10
S2. Activity summary of ivermectin samples in PubChem Alpha-based assays S11
S3. Normalized fluorescence quench rates of ivermectin analogs S14
S4. Compound quality control and source summary S15

Supplementary notes

Note 1. Antiparasitic structure-activity relationships for ivermectin and analogs S16
Note 2. Ivermectin is not an effective SARS-CoV-2 antiviral in high-quality clinical trials S17

Detailed author contributions S18

Supplementary references S19

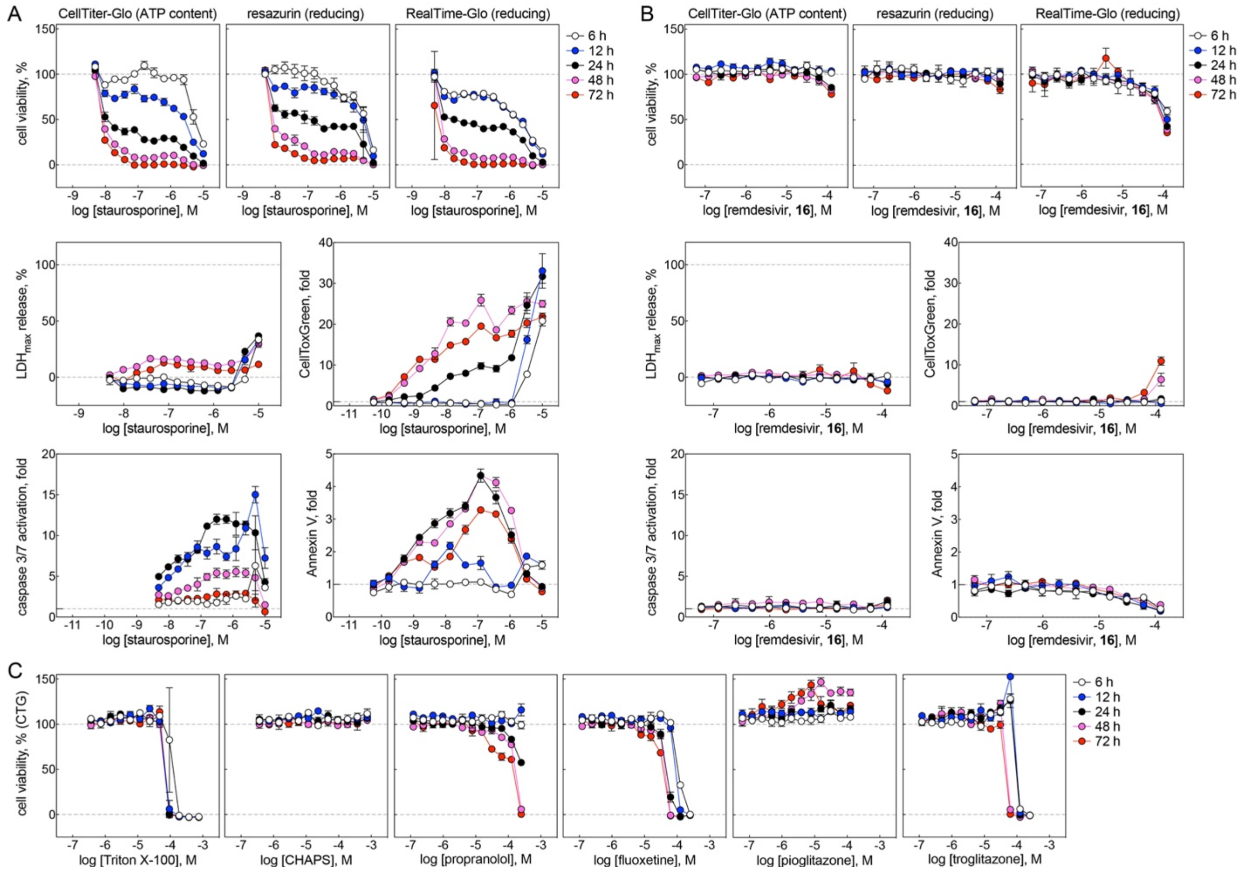


**Supplementary Figure 1. Cellular health control compounds.** (A) Positive cytotoxicity control compound staurosporine decreases cellular viability (CTG, resazurin, RTG), causes A549cytotoxicity (LDH, CellToxGreen), and increases biochemical markers of apoptosis (caspase 3/7, Annexin V) at nanomolar compound concentrations in A549-ACE2 cells. Data are mean ± SD of four intra-plate technical replicates. (B) Control compound remdesivir only decreases cellular viability, causes cytotoxicity, and increases biochemical markers of apoptosis at high micromolar concentrations in A549-ACE2 cells. Data are mean ± SD of four intra-plate technical replicates. (C) Select membrane bilayer modifying control compounds decrease A549-ACE2 viability at micromolar concentrations. Triton X-100, propranolol, fluoxetine, troglitazone: positive cytotoxic controls; CHAPS, pioglitazone: negative cytotoxic controls. Data are mean ± SD of four intra-plate technical replicates. Experiments were performed in parallel with **Figure 2**. Source data are provided as a Source Data file.

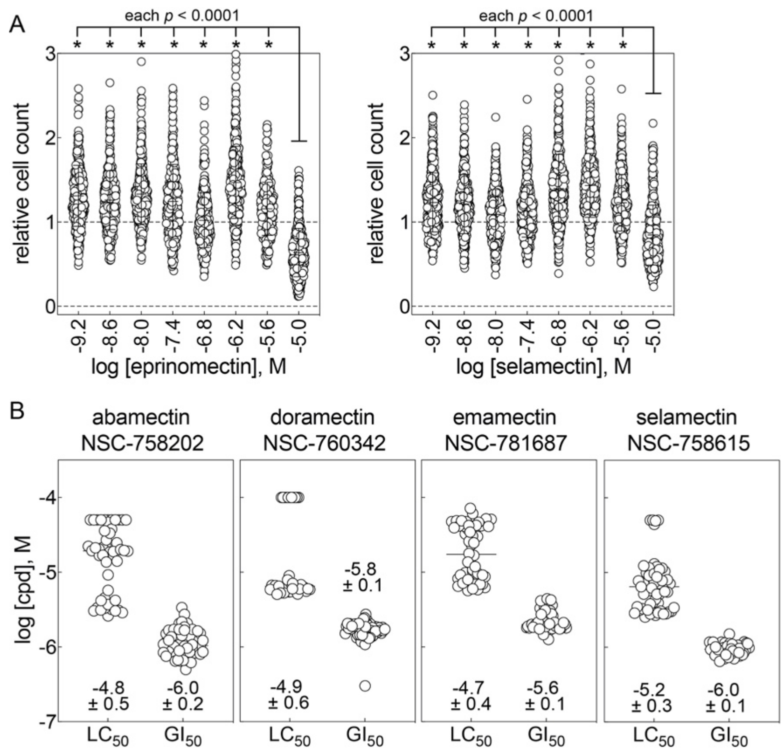


**Supplementary Figure 2. Cellular health ivermectin analogs.** (A) The IVM analogs eprinomectin and selamectin broadly decrease cell counts at low micromolar concentrations in the Broad PRISM assay (Broad DepMap, https://depmap.org/portal/). Each data point represents a unique cancer cell line. Statistical differences determined by ordinary one-way ANOVA adjusted for multiple comparisons. (B) The IVM analogs abamectin, doramectin, emamectin, and selamectin broadly decrease observed cell counts and inhibit cell growth at low micromolar concentrations in the NCI60 assay (https://dtp.cancer.gov). Each data point represents a unique cancer cell line. $LC_{50}$, drug concentration resulting in a 50% reduction in the measured protein at the end of the drug treatment as compared to that at the beginning; $GI_{50}$, drug concentration resulting in a 50% reduction in the net protein increase as measured by sulforhodamine B staining in control cells during the drug incubation. Source data are provided as a Source Data file.

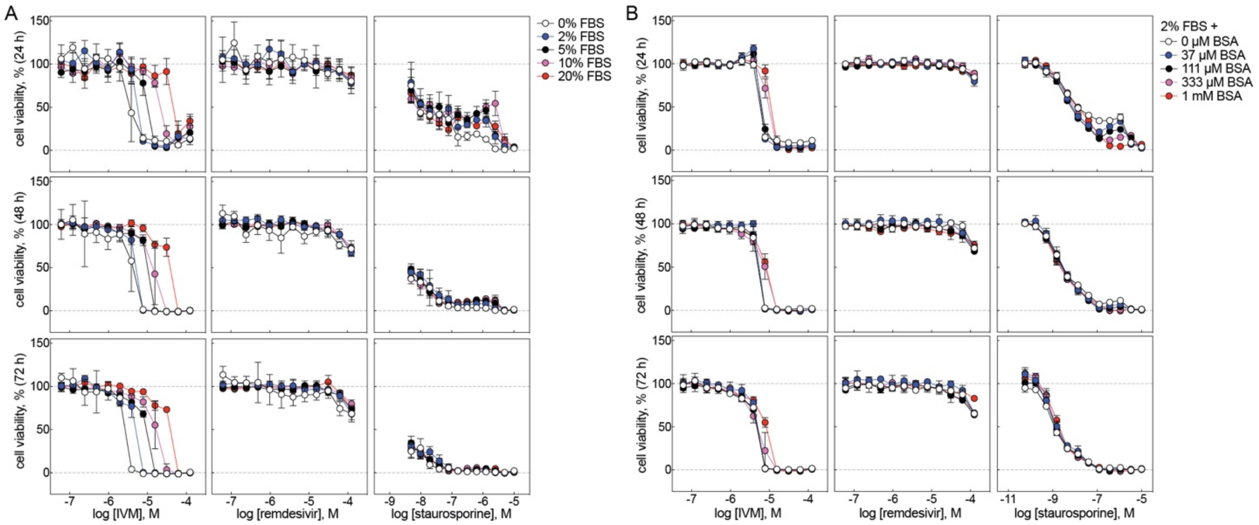


**Supplementary Figure 3. Complete media content can modulate effect of ivermectin on cell viability.** (A) The effect of IVM on A549-ACE2 cell viability depends on the FBS concentration in complete media. This effect is not observed for remdesivir or staurosporine. Data are mean ± SD of four intra-plate technical replicates. (B) The effects of IVM, remdesivir, and staurosporine on A549-ACE2 cell viability does not grossly depend on the BSA concentration in complete media. Note that 2% FBS contains approximately 7.5 μM final albumin concentration, based on an expected albumin concentration of 2.5 g/dL in 100% FBS. Data are mean ± SD of four intra-plate technical replicates. Source data are provided as a Source Data file.

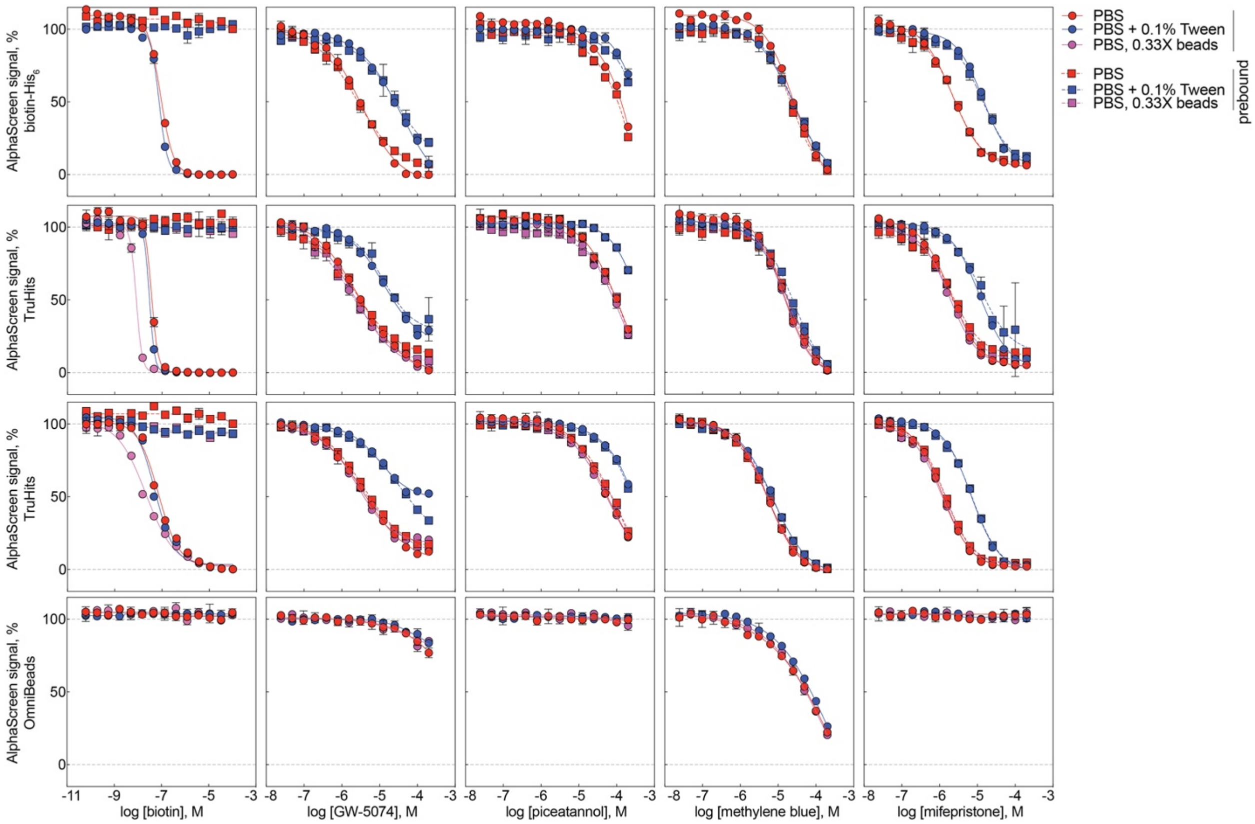


**Supplementary Figure 4. Alpha technology interference profiles of reference compounds.** Compound-mediated interference with the Alpha homogenous proximity technology was assessed according to the NCATS *Assay Guidance Manual*.[1] Biotin, prototypical capture reagent disruptor (streptavidin-biotin interaction); GW-5074, prototypical light scatterer; piceatannol, singlet oxygen quencher; methylene blue, prototypical colored/light absorbance; mifepristone, reported NLS-importin inhibitor.[2] Data are mean ± SD of four intra-plate technical replicates. Source data are provided as a Source Data file.

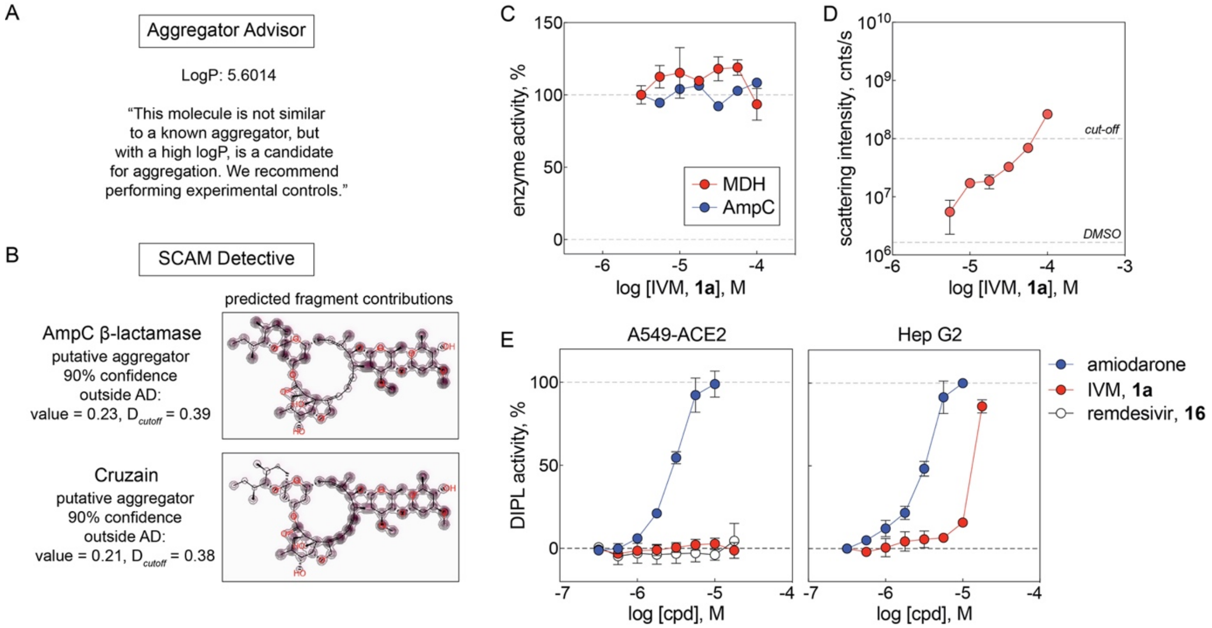


**Supplementary Figure 5. Ivermectin is unlikely to interfere with biological assays by colloidal aggregation or drug-induced phospholipidosis.** (A) IVM is predicted to be a candidate colloidal aggregator by the Aggregator Advisor tool due to its high ClogP (https://advisor.bkslab.org; accessed 07 Oct 2023).[3] Shown is the analysis output. (B) IVM is predicted to be a putative aggregator with high confidence by the SCAM Detective tool (https://scamdetective.mml.unc.edu/; accessed 07 Oct 2023).[4] Shown are the analysis outputs from its models based on AmpC β-lactamase and cruzain, although the structure was flagged as outside the applicability domain. (C) IVM does not inhibit the aggregator-sensitive enzymes AmpC and MDH in the absence of detergent. MDH: data are mean ± SD of three intra-plate technical replicates; AmpC: data are a single point. (D) IVM does not form colloids at low micromolar concentrations as detected by DLS. Data are mean ± SD of three intra-plate technical replicates. (E) IVM does not induce DIPL in A549-ACE2 cells, but induces DIPL at high micromolar concentrations in Hep G2 cells. Amiodarone, positive DIPL control; remdesivir, negative control. Data are mean ± SD of three independent experiments. Amiodarone, positive DIPL control; remdesivir, negative control. Source data are provided as a Source Data file.

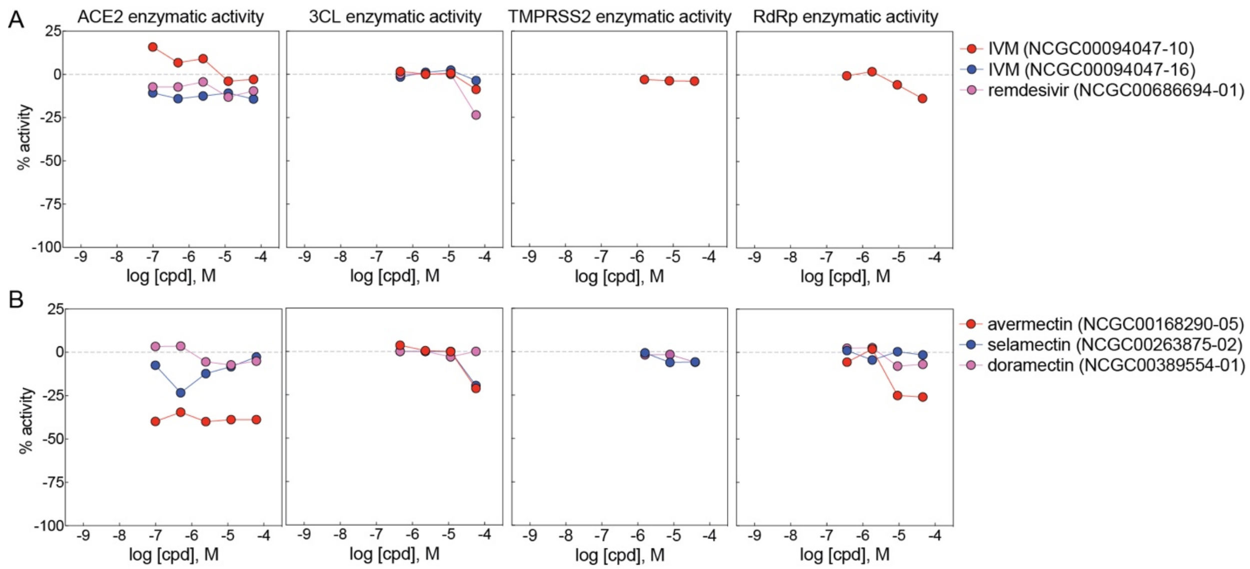


**Supplementary Figure 6. Ivermectin and chemical analogs do not inhibit several SARS-CoV-2-related targets in cell-free assays.** (A) Two IVM samples do not inhibit the SARS-CoV-2 targets 3CL or RdRp, as well as human ACE2 and TMPRSS2, in biochemical assays. Data are single points from an inter-plate qHTS. (B) The IVM analogs avermectin, selamectin, and doramectin do not inhibit the SARS-CoV-2 targets 3CL or RdRp, as well as human ACE2 and TMPRSS2, in biochemical assays. Data are single points from an inter-plate qHTS. Data is from NCATS OpenData Portal. NCGC numbers indicate specific samples. Source data are provided as a Source Data file.

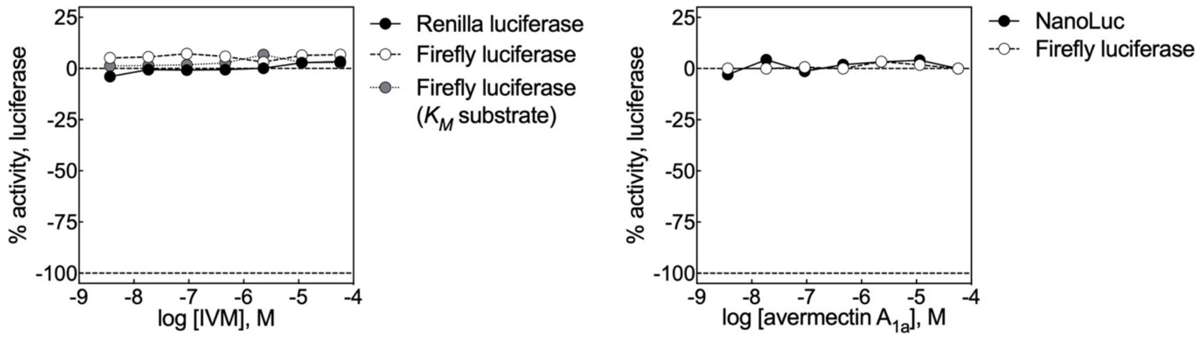


**Supplementary Figure 7. Ivermectin does not inhibit recombinant luciferases.** Data expressed as percent change relative to DMSO control, with 0% being no effect, and -100% being complete inhibition. Data are single points from quantitative high-throughput screening.[5] Source data are provided as a Source Data file.

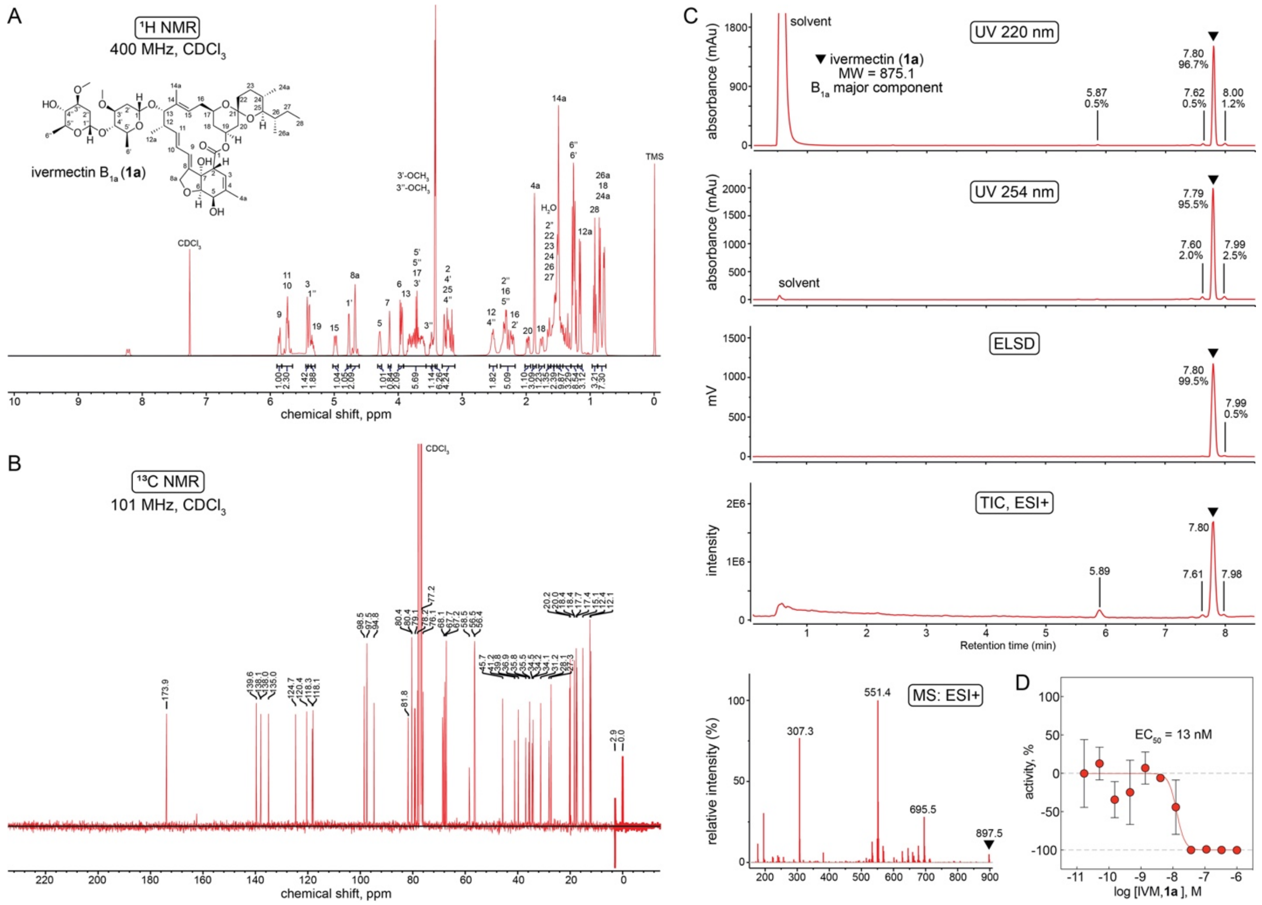


**Supplementary Figure 8. Quality control of primary ivermectin sample.** The primary study sample of IVM in this study (**1a**) was obtained as a USP standard. (A) $^{1}$H NMR of primary IVM sample is consistent with its reported chemical structure.[6] (B) $^{13}$C NMR of primary IVM sample (**1a**) is consistent with its reported chemical structure. (C) Primary IVM sample shows acceptable purity by UPLC-MS, as quantified by UV220 nm, UV254 nm, and ELSD. The parent ion as detected by UPLC-MS (ESI, + mode) is consistent with its reported chemical structure. (D) Primary IVM sample (**1a**) shows expected anthelmintic activity at nanomolar concentrations versus *Caenorhabditis elegans*. Data are mean ± SD of four inter-plate biological replicates. N.b., data included as part of **Figure 1C**. Source data are provided as a Source Data file.

### Supplementary Table 1. Summary of ivermectin target activities.

| target | cys-loop receptor | concentration (range) | reference |
|---|---|---|---|
| GluCl | yes | 100 nM | [7] |
| histamine-gated chloride channel | yes | 1 μM | [8] |
| $H^+$-gated chloride channel | yes | 10 μM | [9] |
| GlyR | yes | 1 μM | [10] |
| $GABA_A$ | yes | 10 μM | [11] |
| $\alpha$7-nAChR | yes | 30 μM | [12] |
| RYR | no | 1 μM | [13] |
| SERCA | no | 10 μM | [13] |
| Na,K-ATPase, H,K-ATPase | no | 10 μM | [14] |
| P2X4 | no | 1 μM | [15] |
| hERG | no | 10 μM | [16*] |
| GIRK | no | 10 μM | [17] |
| various xenobiotic transporters | no | 1 – 10 μM | [18] |

[*], may reflect decreased trafficking[19]

**Supplementary Table 2. Activity summary of ivermectin samples in PubChem Alpha-based assays.** Relevant PubChem biological assays for ivermectin using Alpha-based technology were identified by the following searches: "alphascreen[All Fields] ivermectin", "alphalisa[All Fields] ivermectin", and manual inspection of bioassay records from PubChem CIDs 3085416, 6321424, 11957587, 45114068, and 73265241. Assay activity and protocol information were then extracted by manual review of PubChem assay descriptions and protocols. All assays used AlphaScreen format unless noted. Data accessed 21 July 2023 (https://pubchem.ncbi.nlm.nih.gov/).

| PubChem AID | PubChem CID | PubChem SID | assay name | compound concentration (μM) | activity score | activity summary | detergent | notes |
|---|---|---|---|---|---|---|---|---|
| 1259374 | 11957587 | 321942436 | AlphaScreen-based biochemical high throughput primary assay to identify inhibitors of microphthalmia-associated transcription factor (MITF) | 2.6 | -3.95% | inactive | 0.1% Tween-20 | |
| | | 332950534 | | 2.6 | -33.63% | inactive | | |
| | | 332950174 | | 2.6 | -19.12% | inactive | | |
| | 45114068 | 333494472 | | 2.6 | -3.99% | inactive | | |
| | | 333494081 | | 2.6 | 7.98% | inactive | | |
| 1259310 | 11957587 | 321942436 | AlphaScreen-based biochemical high throughput primary assay to identify activators of the E3 ligase (FBW7) | 26.1 | 0% | inactive | 0.1% Tween-20 | |
| | | 332950534 | | 26.1 | -0.04% | inactive | | |
| | | 332950174 | | 26.1 | 0.43% | inactive | | |
| | 45114068 | 333494472 | | 26.1 | 0.05% | inactive | | |
| | | 333494081 | | 26.1 | 0.12% | inactive | | |
| 1454 | 11957587 | 50106505 | qHTS assay for inhibitors of the ERK signaling pathway using a homogeneous screening assay; stimulation with EGF | 0.460<br>11.49 | -4.79%<br>3.37% | inconclusive | information not provided | lytic cellular assay; lysis buffer may contain detergent |
| 485290 | 11957587 | 50106505 | qHTS assay for inhibitors of tyrosyl-DNA phosphodiesterase (TDP1) | 0.004<br>0.021<br>0.094<br>0.505<br>2.283<br>12.29<br>56.06 | 13.46%<br>4.41%<br>4.11%<br>3.88%<br>3.96%<br>8.38%<br>13.46% | inactive | 0.05% Tween-20 | |
| 1477 | 11957587 | 50106505 | qHTS assay for compounds blocking the interaction between CBF-beta and RUNX1 for the treatment of acute myeloid leukemia | 0.093<br>0.514<br>2.359<br>13.09<br>59.83 | 2.28%<br>5.12%<br>7.18%<br>5.67%<br>0.61% | inactive | 0.01% Tween-20 | |
| 504332 | 11957587 | 50106505 | qHTS assay for inhibitors of histone lysine methyltransferase G9a | 0.018<br>0.092<br>0.501<br>2.335<br>12.70 | 20.01%<br>33.54%<br>32.02%<br>20.37%<br>21.19% | inactive | information not provided | |
| | 73265241 | 90340581 | | 0.004<br>0.018<br>0.092<br>0.501<br>2.335<br>12.70<br>58.95 | -4.98%<br>-6.78%<br>-3.27%<br>-4.71%<br>48.57%<br>33.37%<br>26.62% | inactive | | |

| | | | | | | | | |
|---|---|---|---|---|---|---|---|---|
| 1259251 | 11957587 | 321942436 | AlphaScreen-based biochemical high throughput primary assay to identify inhibitors of Unc-51 like autophagy activating kinase 1 (ULK1) | 5.96 | -0.03% | inactive | 0.01% Triton X-100 | |
| 488949 | 11957587 | 50106505 | qHTS validation assay for inhibitors for MPP8 chromodomain interactions with methylated histone tails | 0.018<br>0.091<br>0.457<br>2.290<br>11.40<br>57.10<br>114.0 | 13.16%<br>-3.34%<br>-3.19%<br>12.13%<br>4.03%<br>-40.23%<br>-39.25% | inconclusive | 0.01% Tween-20 | |
| | 73265241 | 90340581 | | 0.004<br>0.018<br>0.091<br>0.457<br>2.290<br>11.40<br>57.10 | 5.42%<br>9.19%<br>-3.45%<br>-6.88%<br>5.09%<br>-0.41%<br>8.79% | inactive | | |
| 504339 | 3085416 | 56462987 | qHTS Assay for inhibitors of JMJD2A-tudor domain | 0.457<br>2.286<br>11.41<br>57.07<br>113.8 | -9.816%<br>-11.25%<br>-24.22%<br>-28.39%<br>-26.61% | inactive | 0.01% Tween-20 | |
| 720542 | 3085416 | 56462987 | qHTS for inhibitors of AMA1-RON; towards development of antimalarial drug lead: primary screen | 0.464<br>2.32<br>11.61<br>58.24 | -3.38%<br>-6.06%<br>-8.76%<br>-6.41% | inactive | information not provided | |
| 624168 | 3085416 | 56462987 | uHTS identification of small molecule activators of alpha dystroglycan glycosylation | 10 | 0.296% | inactive | 0.01% Tween-20 | cellular assay; AlphaLISA format |
| 504329 | 3085416 | 56462987 | Discovery of small molecule probes for H1N1 influenza NS1A | 12.5 | 9.42% | inactive | none | |
| 623870 | 3085416 | 56462987 | ARNT-TAC3: AlphaScreen HTS to detect disruption of ARNT/TAC3 interactions measured in biochemical system using plate reader - 2158-01_Inhibitor_SinglePoint_HTS_Activity | 9.99 | 0.65% | inactive | 0.02 % Tween-20 | |
| 1272365 | 3085416 | 406214639 | SSB-PriA antibiotic resistant target AlphaScreen | 33 | -2.42% | inactive | 0.01% Triton X-100 | |
| 651724 | 3085416 | 56462987 | qHTS assay for inhibitors of the CtBP/E1A interaction | 57.5 | 7.66% | inactive | 0.02% Tween-20 | |
| 651725 | 3085416 | 56462987 | qHTS assay for inhibitors of the Six1/Eya2 interaction | 57.5 | -2.63% | inactive | 0.02% Tween-20 | |
| 651723<br>651687 | 3085416 | 56462987 | MLPCN PGC1a modulators measured in cell-based system using plate reader - 2139- | 15.56<br>15.56 | -24.50%<br>-9.32% | inactive | information not provided | lytic cellular assay; lysis buffer may contain detergent |

| | | | | | | | | |
|---|---|---|---|---|---|---|---|---|
| | | | 01_Inhibitor_SinglePoint_HTS_Activity | | | | | |
| 540317 | 3085416 | 56462987 | HTS for inhibitors of HP1-beta chromodomain interactions with methylated histone tails | 0.458<br>2.291<br>11.40<br>57.01<br>113.8 | 24.25%<br>20.82%<br>10.03%<br>0.75%<br>7.75% | inactive | 0.01% Tween-20 | |
| 488953 | 73265241 | 90340581 | qHTS validation assay for inhibitors of HP1-beta chromodomain interactions with methylated histone tails | 0.018<br>0.091<br>0.457<br>2.290<br>11.40<br>57.10<br>114.0 | -1.37%<br>-0.01%<br>15.65%<br>15.69%<br>5.59%<br>-7.87%<br>-10.82% | inactive | 0.01% Tween-20 | |
| 743279 | 3085416 | 56462987 | qHTS for inhibitors of inflammasome signaling: IL-1-beta AlphaLISA primary screen | 11.5<br>57.5 | -23.28%<br>-28.88% | inactive | none | nonlytic cellular assay; AlphaLISA format |
| 1259354 | 11957587 | 348438277 | Small-molecule inhibitors of ST2 (IL1RL1) | 17 | information not provided | inactive | Tween-20 | AlphaLISA format |
| 1347059 | 73265241 | 90340581 | CD47-SIRPalpha protein protein interaction – Alpha assay qHTS validation | 0.002<br>0.012<br>0.061<br>0.307<br>1.530<br>7.660<br>38.30 | 1.58%<br>1.24%<br>0.32%<br>1.49%<br>-0.63%<br>-12.51%<br>-12.42% | inactive | 0.05% IGEPAL CA-630 | |

**Supplementary Table 3. Normalized fluorescence quench rates of ivermectin analogs.** Compounds were tested at 10 μM final concentrations. The results in the top eight rows were done using a blinded library; the results in the bottom two rows were controls. Data are mean ± SD for three biological replicates (independent LUV batches). Source data are provided as a Source Data file.

| **compound** | **mean** $\boldsymbol{Rate/Rate_{contorl}}$ | **SD** |
|---|---|---|
| ivermectin, **1a** | 6.5 | 1.3 |
| ivermectin $B_{1a}$ monosaccharide, **3** | 7.7 | 1.3 |
| ivermectin $B_{1a}$ aglycone, **4** | 7.4 | 1.5 |
| doramectin, **8** | 6.1 | 1.5 |
| avermectin $B_{1b}$, **10** | 6.8 | 0.3 |
| delta2-avermectin $B_{1a}$, **11** | 7.2 | 1.1 |
| avermectin $B_{1a}$ aglycone, **12** | 3.6 | 0.5 |
| remdesivir, **16** | 3.4 | 0.4 |
| ivermectin, **1a** | 6.9 | 1.2 |
| 5% EtOH | 5.6 | 1.0 |

### Supplementary Table 4. Compound quality control and source summary.

| cpd | common name | sample ID | vendor | catalog # | ELSD purity, % | UV 220 nm purity, % |
|---|---|---|---|---|---|---|
| **1a** | ivermectin | NCGC00094047-26 | Sigma Aldrich | 1354309 | 99.5 | 96.7 |
| **1b** | ivermectin | NCGC00094047-25 | Santa Cruz Biotechnology | sc-203609 | 98.3 | 94.4 |
| **1c** | ivermectin | NCGC00094047-24 | MedChemExpress | HY-15310 | 99.7 | 95.6 |
| **2** | ivermectin $B_{1b}$ | NCGC00843246-01 | Cayman Chemicals | 23824 | 99.8 | 97.1 |
| **3** | ivermectin $B_{1a}$ aglycone | NCGC00843248-01 | Cayman Chemicals | 19442 | 99.9 | 97.7 |
| **4** | ivermectin $B_{1a}$ monosaccharide | NCGC00843239-01 | Cayman Chemicals | 19443 | 99.7 | 95.1 |
| **5** | 2,3-dehydro-3,4-dihydro ivermectin | NCGC00843242-01 | Cayman Chemicals | 23849 | 99.6 | 96.7 |
| **6** | 28-oxo ivermectin $B_{1a}$ | NCGC00843250-01 | Toronto Research Chemicals | O856970 | 99.4 | 96.3 |
| **7** | epi-ivermectin $B_{1a}$ | NCGC00843241-01 | Cayman Chemicals | 25211 | 99.4 | 96.2 |
| **8** | doramectin | NCGC00390529-07 | Cayman Chemicals | 19467 | 99.7 | 96.8 |
| **9** | selamectin | NCGC00095066-06 | Cayman Chemicals | 21529 | pass | pass |
| **10** | avermectin $B_{1b}$ | NCGC00843247-01 | Cayman Chemicals | 17453 | 93.4 | 86.8 |
| **11** | delta2-avermectin $B_{1a}$ | NCGC00843243-01 | Cayman Chemicals | 25119 | 99.4 | 94.6 |
| **12** | avermectin $B_{1a}$ aglycone | NCGC00843245-01 | Cayman Chemicals | 28051 | 99.4 | 95.0 |
| **13** | avermectin $B_{1a}$ monosaccharide | NCGC00843244-01 | Cayman Chemicals | 26690 | 94.2 | 87.0 |
| **14** | abamectin | NCGC00168290-11 | Cayman Chemicals | 19201 | - - | 89.4 (mixture) [1] |
| **15** | moxidectin | NCGC00163732-08 | MedChemExpress | HY-B0777 | - - | 94.4 [1] |
| **16** | remdesivir | | Cayman Chemicals | 30354 | pass | pass |

[1], UV 254 nm purity

**Supplementary Note 1. Antiparasitic structure-activity relationships for ivermectin and analogs.** The structure-activity relationships (SAR) of ivermectin and avermectin analogs has been extensively studied in a wide variety of anthelmintic, insecticidal, and acardicidal models.[20,21] Interpreting SAR for these compounds can be complicated given their broad spectrum of activity against helminths and arthropods. However, there is multiple lines of evidence that minor structural modifications can cause significant changes in biological activity which span several orders of magnitude in terms of compound concentration. In models of *Tetranychus urticae* (red spider mite), there is a greater than ten-fold change in activity between the disaccharide and monosaccharides ivermectins/avermectins compared to their corresponding aglycones.[20] Similar findings showing the importance of the sugar moieties have been reported in other systems.[21] When tested in a *Haemonchus contortus* larval development assay, ivermectin and avermectin analogs showed evidence of SAR, including activity enhancement with hydroxylation (vs. oxo and oxime groups) at the C-5 position, and activity enhancement with double bonds at the C-22/C-23 positions in combination with a *sec*-butyl/isopropyl substituent at the C-25 position.[22] The importance of the hydroxyl group at the C-5 position was also shown in a *Trichostrongylus colubriformis* model.[23] Related, a study focusing on the inhibitory ability of avermectin analogs on *Tetranychus cinnabarinus* (carmine spider mite) showed SAR spanning four orders of magnitude.[24] Some modifications do not appear to lead to significant changes in activity (using a *Trichostrongylus colubriformis* model), however, such as acylation at the C-4'' position.[23]

**Supplementary Note 2. Ivermectin is not an effective SARS-CoV-2 antiviral in high-quality clinical trials.** Several reports have shown a lack of clinical efficacy of IVM for the treatment of SARS-CoV-2 in high-quality (e.g., double-blind, randomized, placebo-controlled) clinical trials.[25, 26, 27, 28, 29, 30]

**Detailed author contributions**

Performed Alpha technology counter-screens: JLD. Performed analytical chemistry studies (NMR, UPLC-MS): SR. Performed *C. elegans* assays: PD. Performed cellular health assays: JLD. Performed drug-induced phospholipidosis counter-screens: ADW. Performed GPCR profiling assays: XPH, BLR. Performed membrane bilayer studies: RR. Performed PubChem analyses: JLD. Performed voltage-clamp studies on voltage-dependent sodium channels: KFH. Performed SARS-CoV-2 live virus assays: RTE. Performed SARS-CoV-2 high-content assay data analyses: TCV.
