## Supplementary material for "Nonspecific membrane bilayer perturbations by ivermectin underlie SARS-CoV-2 *in vitro* activity": Source Data 2: EZX_smpl.INI

```
[HEADER]
SUMFILE=
JOBID=11-29-21 14-42-32.272
NAME=NCGC00094047-26
INFO=0380650810
LOGINDATE=11/29/21 14:42:33
TRAYTYPE=Multisampler Tray
PLATETYPE=*54VialPlate*
SUBMISSIONUNIT=2mL vials
POSITION=D1F-A4
SAMPLES=4
USER=ZII
GROUP=WalkUp Chemist
[MS_VB_VARS]
TARGET_1=874.51
MOLFORM_1=
CUSTOM_1=FINAL GRD BUT NO PRINT
CUSTOM_2=0380650810
CUSTOM_3=FINAL_GRD_NO PRINT
CUSTOM_1_DISPNAME=WalkUp Method Description
CUSTOM_2_DISPNAME=WalkUp Sample Description
CUSTOM_3_DISPNAME=WalkUp Method
[ASR]
EZX_NoUserSetUVWavelength=1
EZX_UserUVWavelength=
```
