## Supplementary material for "Nonspecific membrane bilayer perturbations by ivermectin underlie SARS-CoV-2 *in vitro* activity": Source Data 2: Report.PDF

**File ..mschem\11-21\291121-NCGC00094047-26-03510.D Tgt Mass (EZX): 874.51**
**Injection Date : 29-Nov-21, 15:14:33 Seq. Line : 0**
**Sample Name : NCGC00094047-26 Location : D1F-A4**
**Acq. Operator : Zina Itkin Inj : 1**
**Spec. Reported : UV Integration Inj Volume : -3 ul**
**Acq. Method : C:\Users\Public\Documents\ChemStation\1\Methods\FINAL_GRAD_NO_PRINT.M**
**Analysis Method : C:\Users\Public\Documents\ChemStation\1\Methods\FINAL_GRAD_NO_PRINT.M**
**Sample Info : 0380650810 WalkUp method: 'FINAL_GRD_NO PRINT' Mol Wt: 874.51**
**Method Info : FINAL GRD BUT NO PRINT**

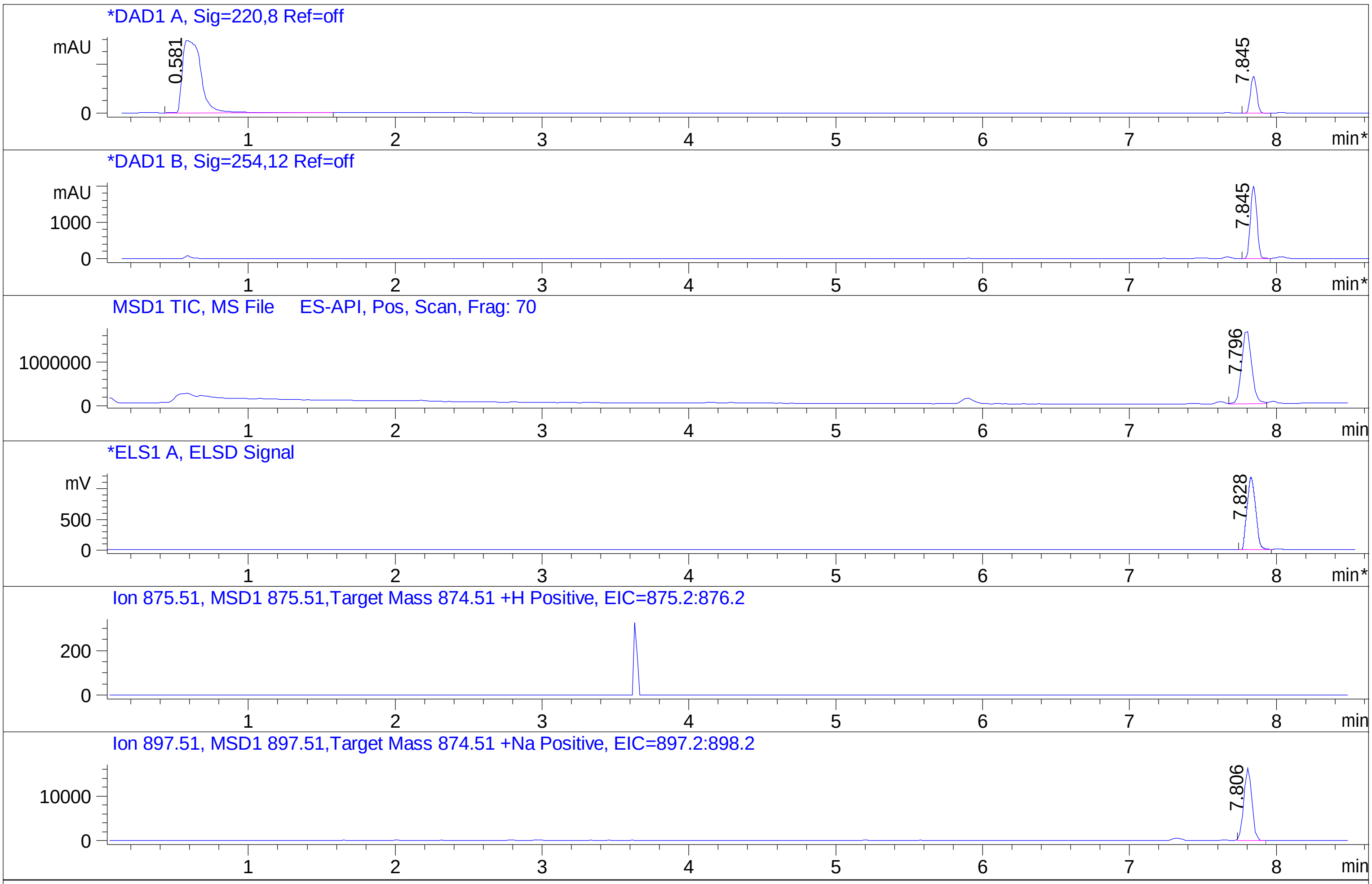


Integration Results for DAD1 A, Sig=220,8 Ref=off

| RetTim | Width | Area | Height | Area% | MS(+) |
|---|---|---|---|---|---|
| 0.58 | 0.12 | 25976.43 | 2953.10 | 85.49 | 179 |
| 7.84 | 0.05 | 4410.07 | 1522.47 | 14.51 | 551 |

Integration Results for DAD1 B, Sig=254,12 Ref=off

| RetTim | Width | Area | Height | Area% | MS(+) |
|---|---|---|---|---|---|
| 7.85 | 0.05 | 6051.58 | 2001.91 | 100.00 | 551 |

Ret. Time: 0.58  <<<< POSITIVE SPECTRA >>>>  Ret. Time: 7.84

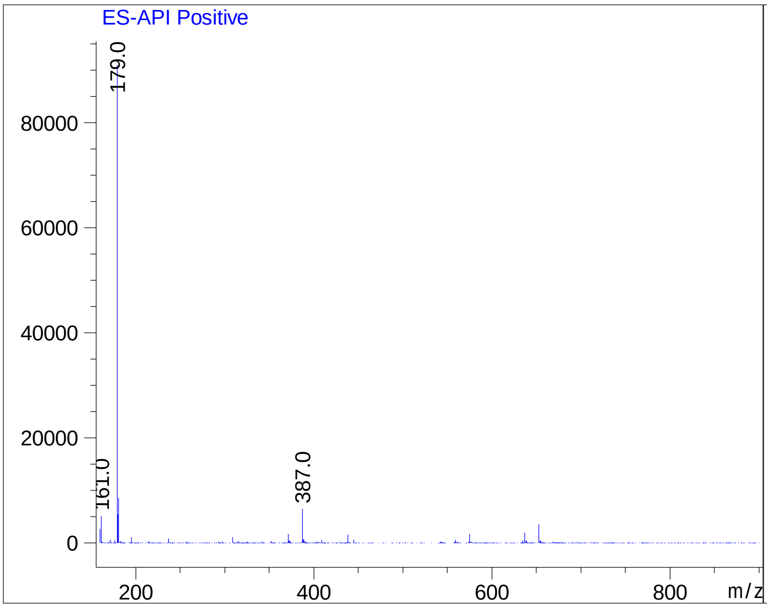


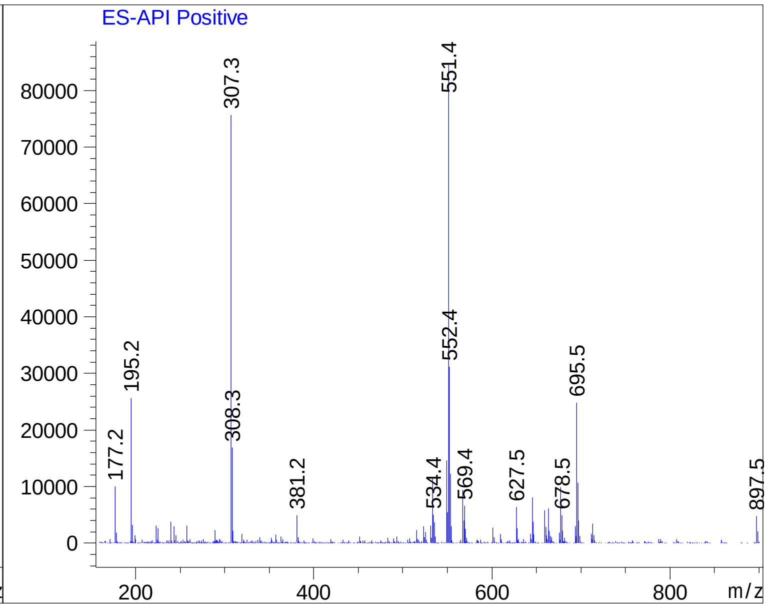
